# Individual strategies for ignoring irrelevant information are reflected in distinct neural signatures during temporal attention

**DOI:** 10.64898/2026.06.23.733825

**Authors:** M. Gironimi, K.I. Ryom, T. Potracov, A. Orsini, F. Pulecchi, M.E. Diamond

## Abstract

Attentional functions enable the nervous system to regulate incoming information, selecting what is behaviorally relevant while filtering out distractions. To investigate attentional control in rats, we developed a paradigm in which animals are required to ignore an irrelevant tactile stimulus (a vibration) and categorize the relevant one as weak or strong. Stimuli were separated in time but delivered to the same set of vibrissae, making this a temporal rather than spatial attention task. In the first task version, the irrelevant stimulus was presented first and the relevant second. Across animals, we observed substantial variability in how this task was learned and performed, consistent with emerging work showing that learning unfolds along idiosyncratic trajectories rather than converging on a single behavioral solution. While the irrelevant stimulus typically exerted an attractive bias on judgments, some rats progressively reduced this influence and achieved near-complete suppression, transitioning from a proficient to an expert stage of performance. Other animals, however, adopted alternative stable strategies and did not exhibit comparable levels of attentional filtering.

To probe behavioral flexibility, we designed a version of the task in which the relevant stimulus could appear in either temporal position. Under these conditions, rats generally struggled to flexibly allocate attention across time, again with substantial variability across individuals. Importantly, behavioral analyses indicate that this variability is not random but reflects a small number of reproducible strategy classes, characterized by distinct patterns of sensitivity to stimulus order. These findings suggest that attentional control in this task does not rely on a single canonical algorithm, but instead emerges from a constrained set of alternative solutions.

Electrophysiological recordings from a limited number of animals revealed neural correlates consistent with these behavioral differences. In expert animals performing the original task, neuronal populations in motor cortex showed differential encoding of the irrelevant and relevant stimuli. In a proficient animal implanted in both vS1 and M1/M2, outcome-dependent modulation was prominent in vS1, whereas M1/M2 transformed sensory inputs into categorical representations with partial suppression of the irrelevant stimulus. Local field potential analyses further indicated that effective task performance was associated with increased high-gamma synchronization between vS1 and M1/M2, alongside low-beta modulation consistent with top-down interactions.

Taken together, these results suggest that learning to ignore irrelevant information reshapes interactions between sensory and motor cortices, but that this process unfolds heterogeneously across individuals. Rather than reflecting a single mechanism of attentional control, the data support a framework in which multiple, identifiable strategies coexist within a shared task structure, underscoring the importance of individual differences in understanding the neural basis of attention.

## Introduction

We are constantly flooded with streams of sensory information. Executive functions enable goal-directed behavior, including the ability to ignore stimuli that are irrelevant to current objectives and to selectively attend to those that are informative. While visuospatial attention has been extensively studied in humans and non-human primates [1–4], less is known about how rodents allocate attention within a single sensory modality.

In the current study, we investigate how rats learn to ignore irrelevant vibrotactile information, focusing on both the behavioral and neural dynamics of temporal attention. In our paradigm, the relevant and irrelevant stimuli were physically indistinguishable; as a result, distinguishing relevant from irrelevant inputs was therefore not an innate capacity but an acquired one, requiring prolonged training.

This intensive, longitudinal interaction allowed us to ask whether animals converged on a single solution to the task or instead adopted individual-specific strategies. More generally, recent work has emphasized that learning and behavior do not necessarily converge onto a single canonical solution, but instead unfold along individual-specific trajectories, with different animals adopting distinct but internally consistent strategies [5, 6]. We therefore asked whether attentional control in this paradigm would be homogeneous or heterogeneous across animals.

To address these questions, we developed the Irrelevant Stimulus Paradigm, a task in which two vibrotactile stimuli are delivered sequentially, separated by a variable inter-stimulus interval. In the first version of the task, the relevant stimulus is tagged by an acoustic cue and always appears in the second temporal position. Rats are required to ignore the first, non-tagged stimulus and report the intensity of the relevant one. By delivering both stimuli to the same set of vibrissae, the task isolates temporal attention rather than spatial selection.

This design allowed us to examine how the irrelevant stimulus biases judgments about the relevant one, as well as how such biases are modulated through training. Because the relevant stimulus always appeared in the second temporal position in the standard task, animals could, in principle, rely on stimulus order rather than attentional selection. To control for this possibility, we included tests in which the acoustic cue was removed, allowing us to dissociate the contribution of temporal position and cue-based selection.

To further assess behavioral flexibility, we introduced a second version of the task in which the relevant stimulus could occur in either temporal position, forcing animals to rely exclusively on the acoustic cue. This manipulation enabled us to test whether rats can flexibly allocate attention across time, rather than adopting fixed strategies tied to stimulus order.

In parallel, we investigated how sensory and motor cortical areas represent relevant and irrelevant information and how their interaction evolves with learning and expertise. Electrophysiological recordings were obtained in rats performing both the standard and dynamic tasks at different stages of training.

Frontal areas are known to exert top-down influences on early sensory cortex, shaping activity to enhance the processing of behaviorally relevant stimuli [7–10]. Here, we asked whether similar mechanisms operate in the rodent somatosensory system during temporal attention. Using population decoding and local field potential analyses, we characterized (i) how relevant and irrelevant stimuli are represented in vS1 and M1/M2, (ii) how these representations depend on behavioral performance and training stage, and (iii) how interactions between these areas support stimulus selection.

## Results

### Irrelevant stimuli systematically bias perceptual judgments

#### Effects of irrelevant stimuli on perceptual judgments

We investigated whether rats can ignore task-irrelevant sensory inputs when forming perceptual judgments. To this end, we developed the Irrelevant Stimulus Paradigm (ISP), in which animals judged the intensity of a vibrotactile target stimulus (REL) while ignoring a temporally separated, non-cued distractor stimulus (IRR). In the standard version of the task, the relevant stimulus always occurred in the second temporal position (Figure 1).

**Figure 1:**
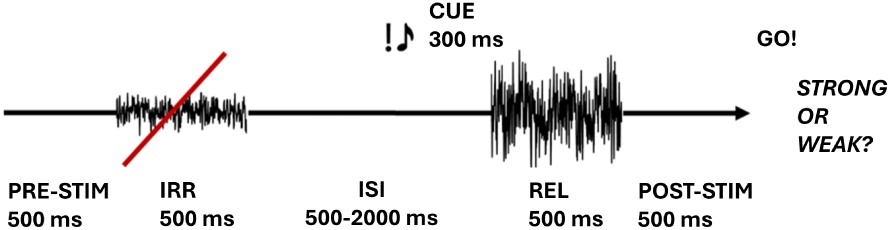
Irrelevant Stimulus Paradigm. The first variant of the *Irrelevant Stimulus Paradigm*, called standard task.

Each trial began with nose-poke fixation, followed by presentation of IRR, a variable inter-stimulus interval (ISI; 500–2000 ms), and then REL, whose onset was signaled by an acoustic cue. Rats reported whether REL was “strong” or “weak” via their choice to lick at the L or R spout. Stimulus intensity was defined as the nominal mean velocity of a sequence of vibration samples drawn from Gaussian distributions, yielding nine discrete intensity levels for both REL and IRR.

Approximately 20% of the trials within each training session was constituted by *single-stimulus* trials, in which only the relevant stimulus was delivered. These trials were introduced to ensure that rats remained in the nosepoke during presentation of the first stimulus, and to exclude the possibility that they remained in the nose poke and ignored events for a fixed period of time (see Supplementary Material).

All rats acquired stable performance, as indicated by steep psychometric functions (Figure 2, top), consistent with the use of reliable stimulus–response strategies. However, performance was systematically biased by the irrelevant stimulus. When conditioned on IRR intensity, psychometric functions shifted toward the category of the distractor, indicating an attractive bias: REL was more likely to be judged “strong” when preceded by a strong IRR, and “weak” when preceded by a weak IRR (Figure 2).

**Figure 2:**
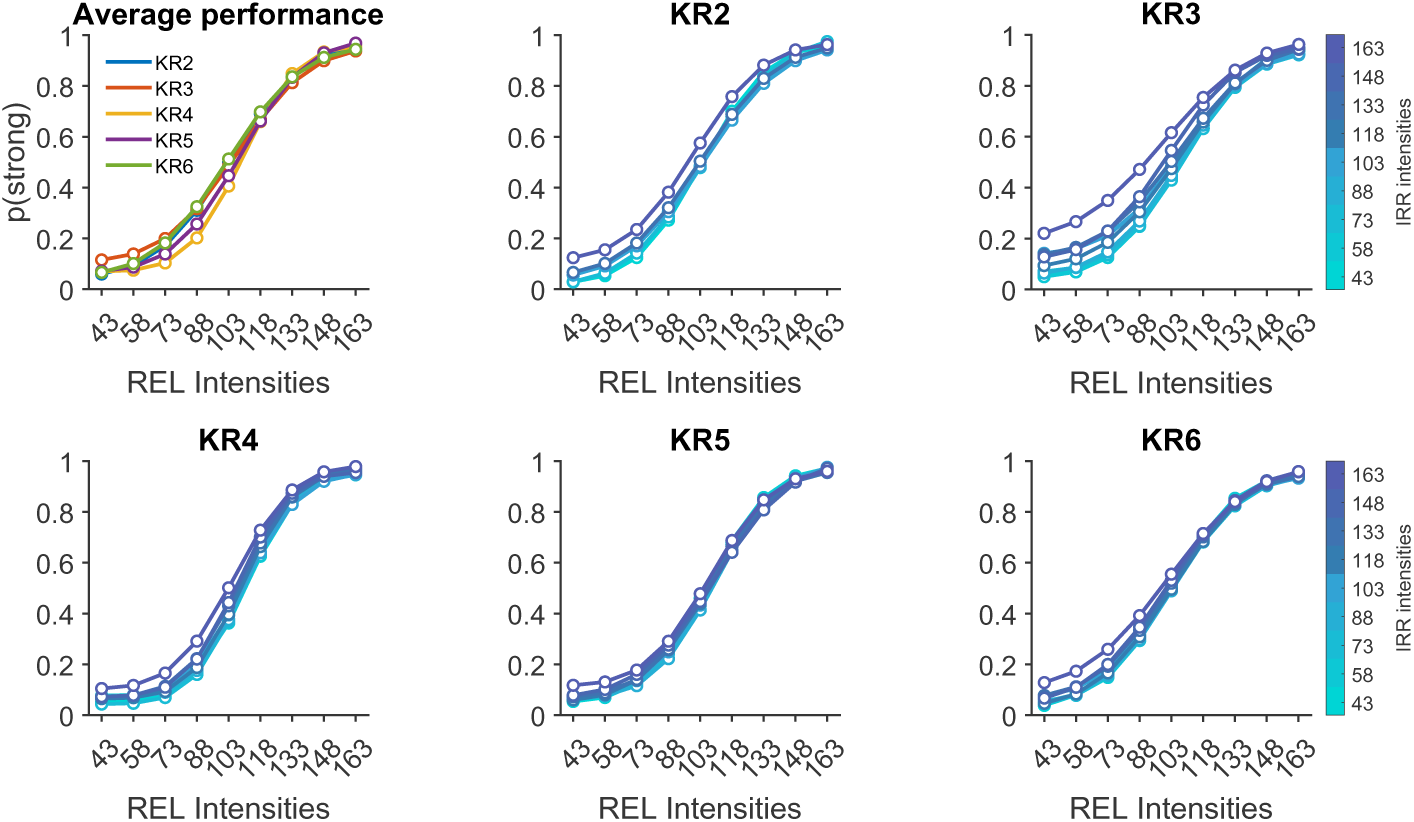
Average performance and IRR bias, for individual rats. Average behavioral performance for individual rats (top left), and IRR induced bias. Individual rats exhibited varying degrees of IRR bias’ magnitude. Dots represent fitted probabilities at each stimulus intensity, and lines connect consecutive fitted probabilities.

This effect showed two notable features. First, the bias was asymmetric, primarily affecting weak REL stimuli, whereas strong REL stimuli were largely resistant to interference. Second, its magnitude varied substantially across individuals, ranging from negligible (e.g. KR5) to pronounced (e.g. KR3), indicating marked inter-individual variability.

The temporal separation between IRR and REL modulated the strength of this bias. Short ISIs enhanced the effect of strong IRR stimuli, whereas longer intervals reduced it (Figure 3b). In contrast, the influence of weak IRR stimuli was more heterogeneous across rats, with some individuals showing little or no dependence on ISI duration (Figure 3a).

**Figure 3:**
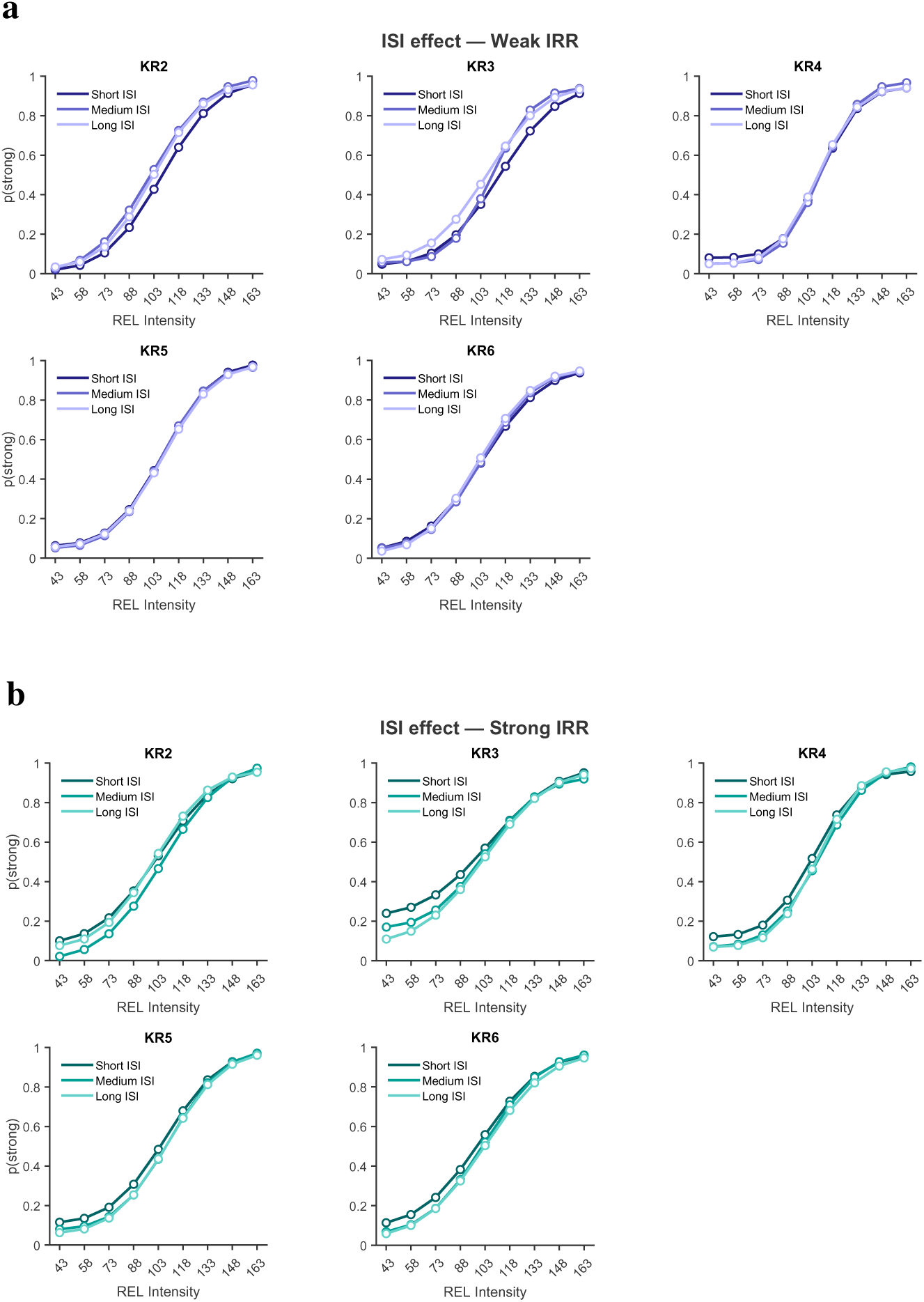
Effect of the ISI duration on the IRR bias, for individual rats. **a.** Effect of ISI duration when IRR was weak. **b.** Effect of ISI duration when IRR was strong.

### Rats rely on temporal structure to infer stimulus relevance

#### Effect of cue removal on performance

In the standard task, the relevant stimulus was always presented second. This raises the possibility that rats used temporal position, rather than the acoustic cue, to determine stimulus relevance. To test this, we removed the acoustic cue in a modified version of the task (Figure 4) and compared performance before and after cue removal. In this version, approximately 40% of trials were constituted by single-stimulus trials (see Supplementary Material).

**Figure 4:**
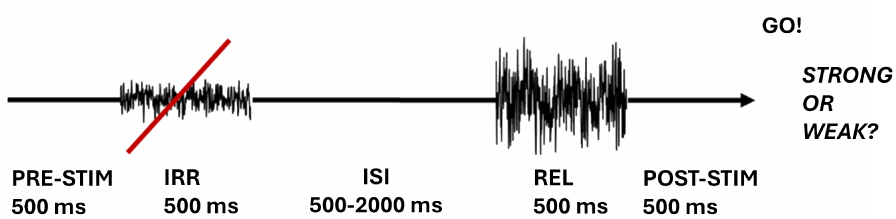
ISP modification: control task. Schematization of the control task, used to test whether rats also used the acoustic cue to assess relevance.

Psychometric functions remained largely unchanged following removal of the cue, with the exception of one subject (KR4), which showed a moderate decrease in performance (Figure 5). These results indicate that rats primarily relied on the temporal structure of the stimulus sequence to infer relevance, while the acoustic cue contributed only marginally.

**Figure 5:**
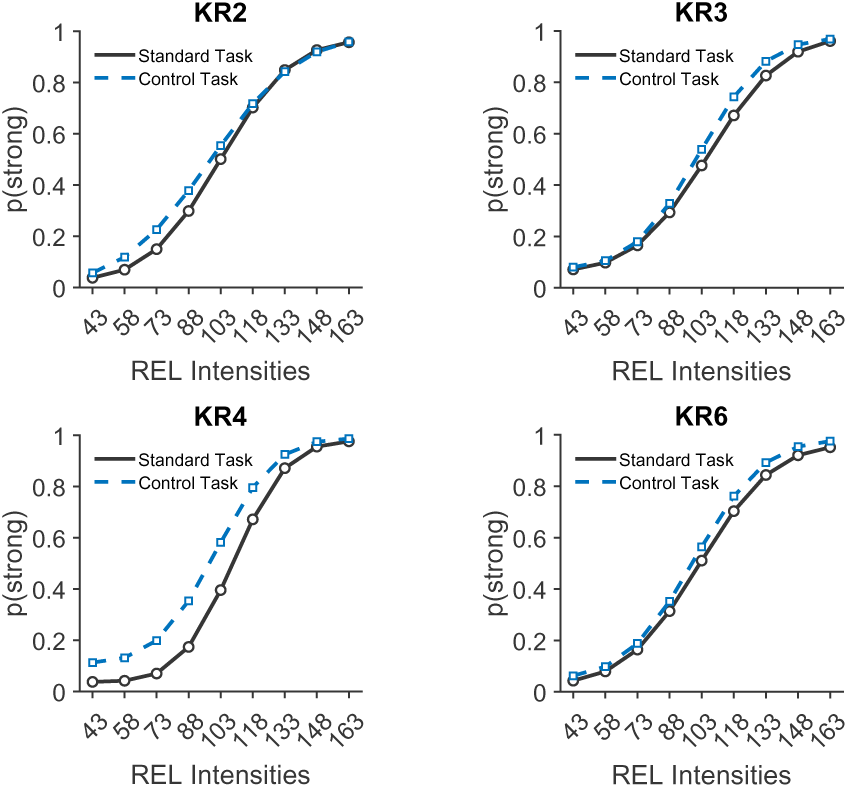
Average performance during the standard task and control task. Comparison of the average performance during the standard task and control task, for individual subjects.

### Limited flexibility in allocating attention across time

#### Performance in the dynamic task

We next asked whether rats could flexibly allocate attention when stimulus relevance varied across trials. In a further variant of the ISP, the position of the relevant stimulus was randomized within each session: half of the trials contained REL in the second position (IRR–REL), and half in the first position (REL–IRR) (Figure 6). Animals trained in this task variant, were exposed to REL-IRR trials and IRR-REL trials since the early training phases.

**Figure 6:**
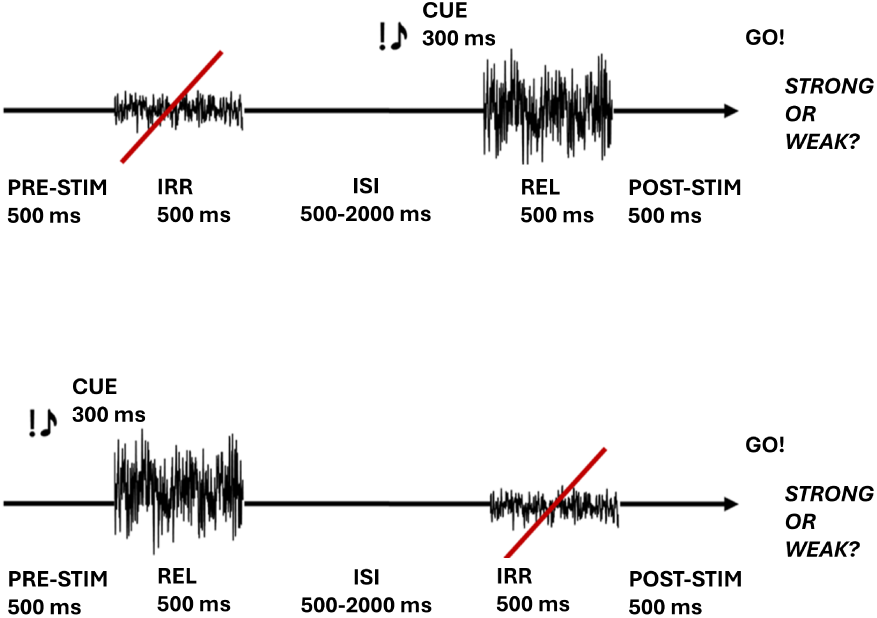
ISP modification: dynamic task. Schematization of the dynamic task, used to test the attentional flexibility of rats across time.

Across all animals, performance was higher when the relevant stimulus appeared in the second position (IRR–REL trials) than when it appeared first (REL–IRR trials) (Figure 7). The attractive bias induced by IRR was present in both trial types, but was stronger when REL occurred first (Figure 8), suggesting a tendency to overweight later stimuli in the sequence.

**Figure 7:**
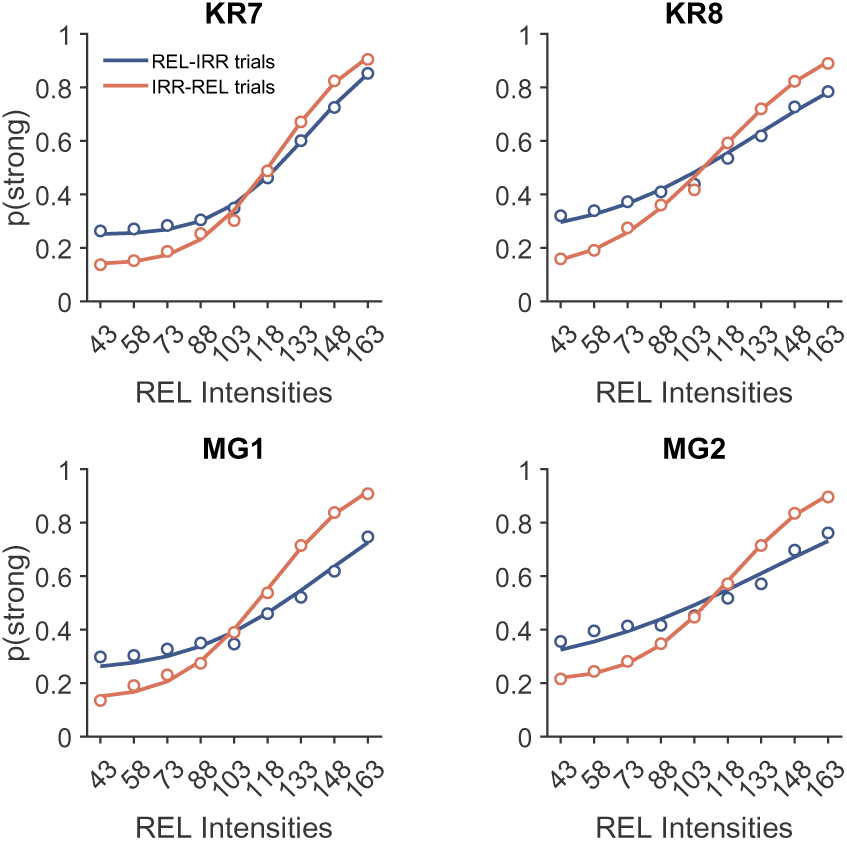
Average performance during the dynamic task. Psychometric functions showing the average performance for each trial type, for single subjects.

**Figure 8:**
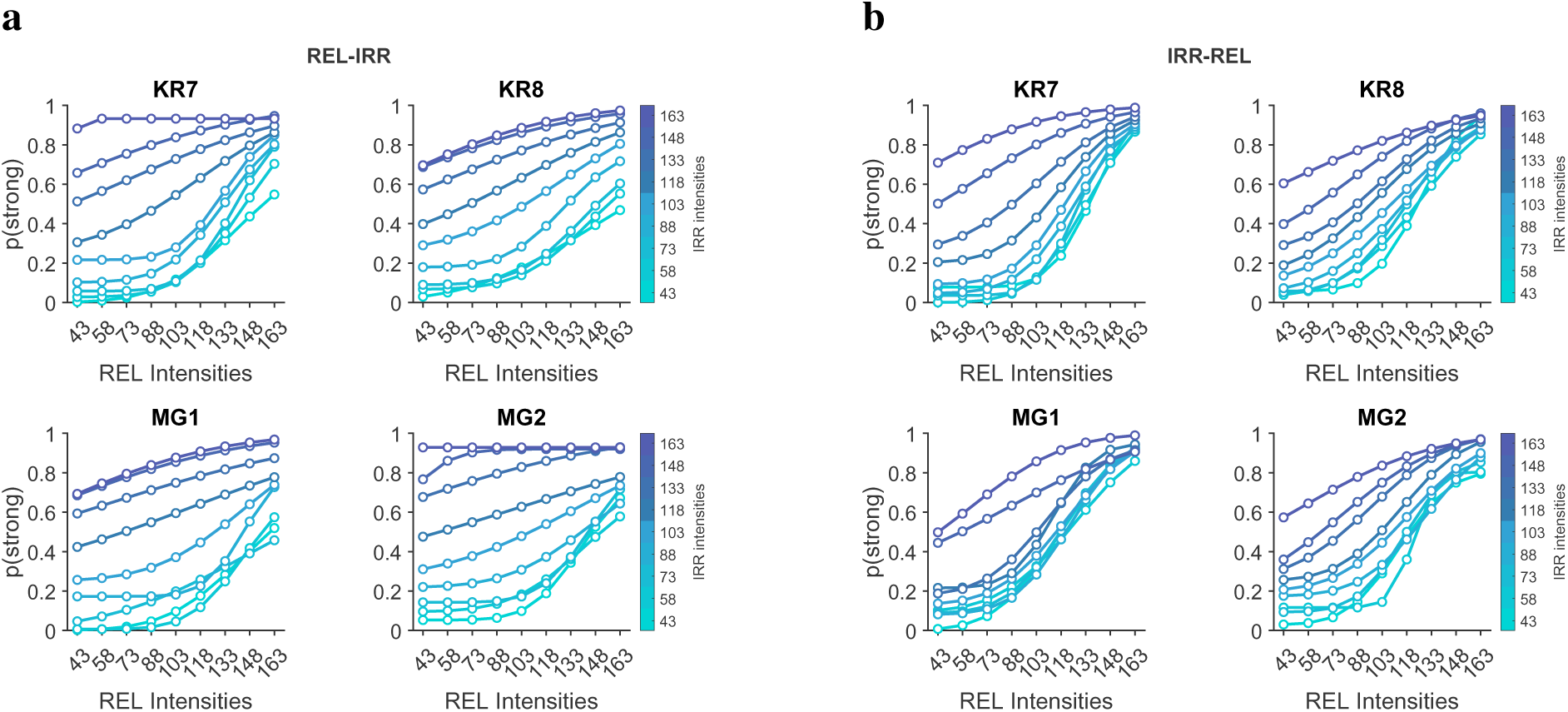
Psychometric functions describing the IRR effect in the dynamic task. **a.** Psychometric functions showing the IRR-induced bias for REL-IRR trials, for single subjects. **b.** Same, for IRR-REL trials.

#### Evidence for a second-stimulus heuristic

To test this directly, we recomputed psychometric functions by plotting choice as a function of IRR rather than REL. Now, performance improved selectively in conditions in which the IRR in the second position as compared to IRR in the first position (Figure 9), consistent with the use of a simple heuristic favoring the use of the second stimulus, irrespective of its actual relevance according to the task rule.

**Figure 9:**
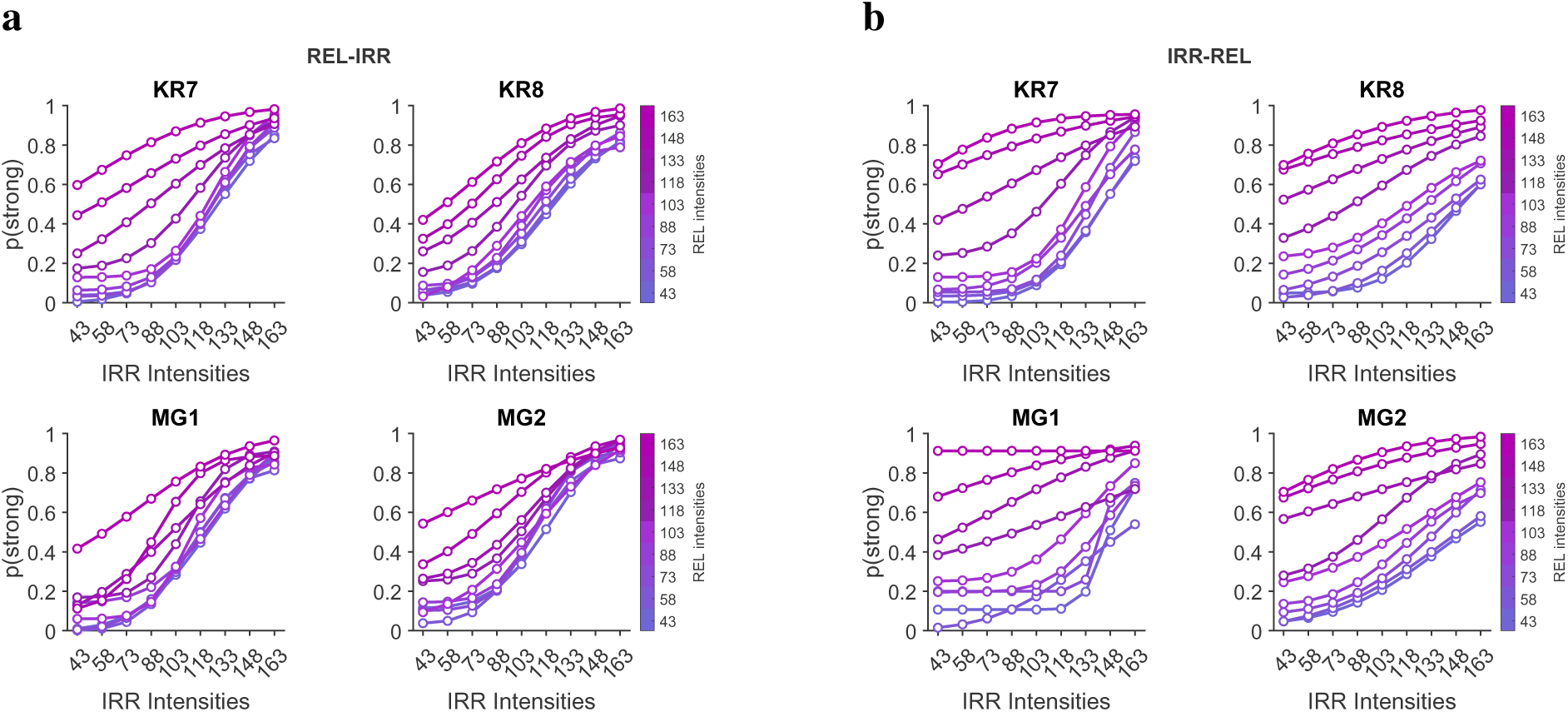
Psychometric functions assuming rats judge IRR as the relevant stimulus. **a.** Psychometric functions showing the IRR-induced bias for REL-IRR trials, for single subjects. **b.** Same, for IRR-REL trials.

### Learning reduces the influence of irrelevant stimuli when task structure is stable

We next examined how sensitivity to irrelevant information evolved with training, relative to the standard task. For each subject, we estimated the contribution of REL and IRR to choice using a generalized linear model (GLM) applied to successive bins of training sessions.

Rats exhibited distinct learning trajectories. In some animals (KR2, KR3, KR4), the weight assigned to IRR progressively decreased over training, approaching zero, consistent with a gradual suppression of irrelevant information (Figure 10). In others (KR5, KR6), IRR weights were near zero from early in training, indicating rapid acquisition of an effective strategy.

**Figure 10:**
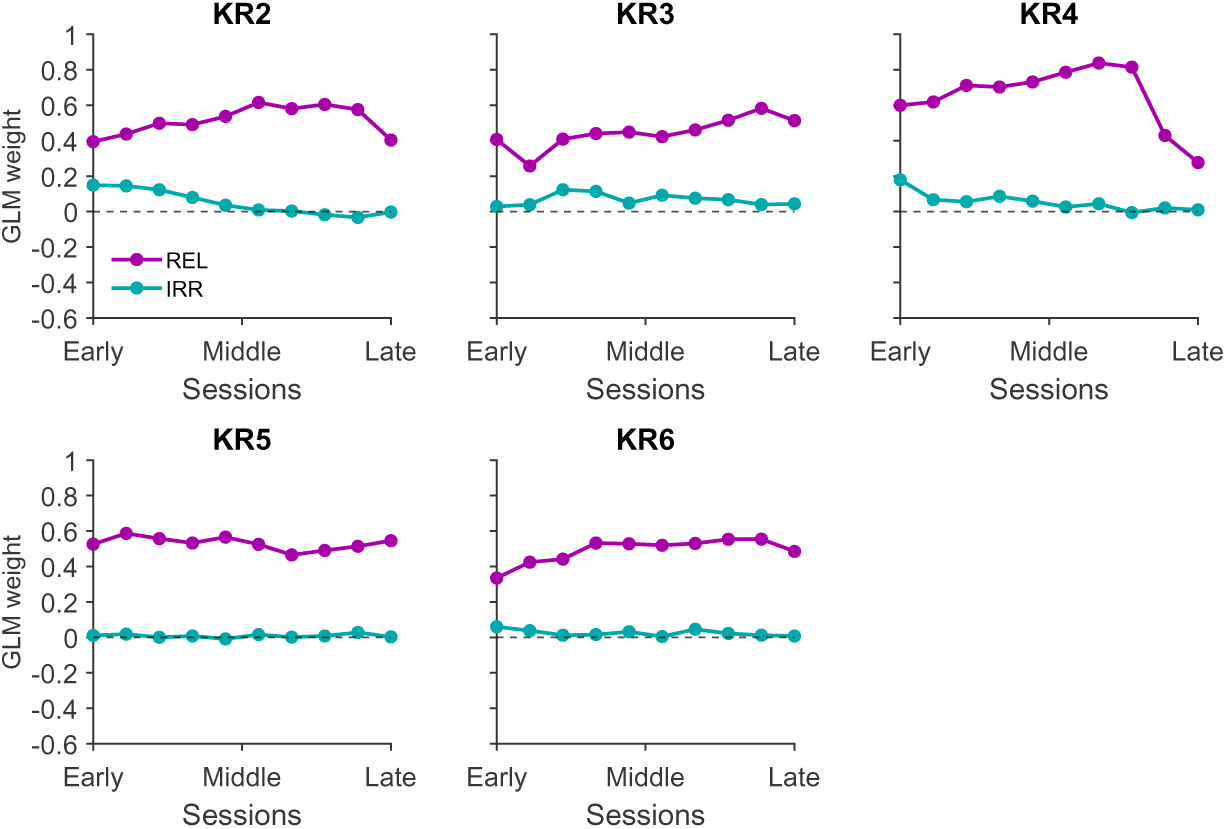
Progression of learning (standard task). Contribution of REL and IRR stimuli to the choice, as described by the GLM weights. All training sessions were binned in 10 groups, and a GLM was computed for each group.

Based on these dynamics, we operationally defined two behavioral regimes: a proficient stage, in which rats performed the task successfully but remained susceptible to IRR-induced bias, and an expert stage, in which IRR exerted little to no influence on behavior. In rats showing gradual learning, the transition between these stages was accompanied by a clear reduction in bias magnitude (Figure 11), whereas in rapidly learning rats no such transition was apparent (Figure 12).

**Figure 11:**
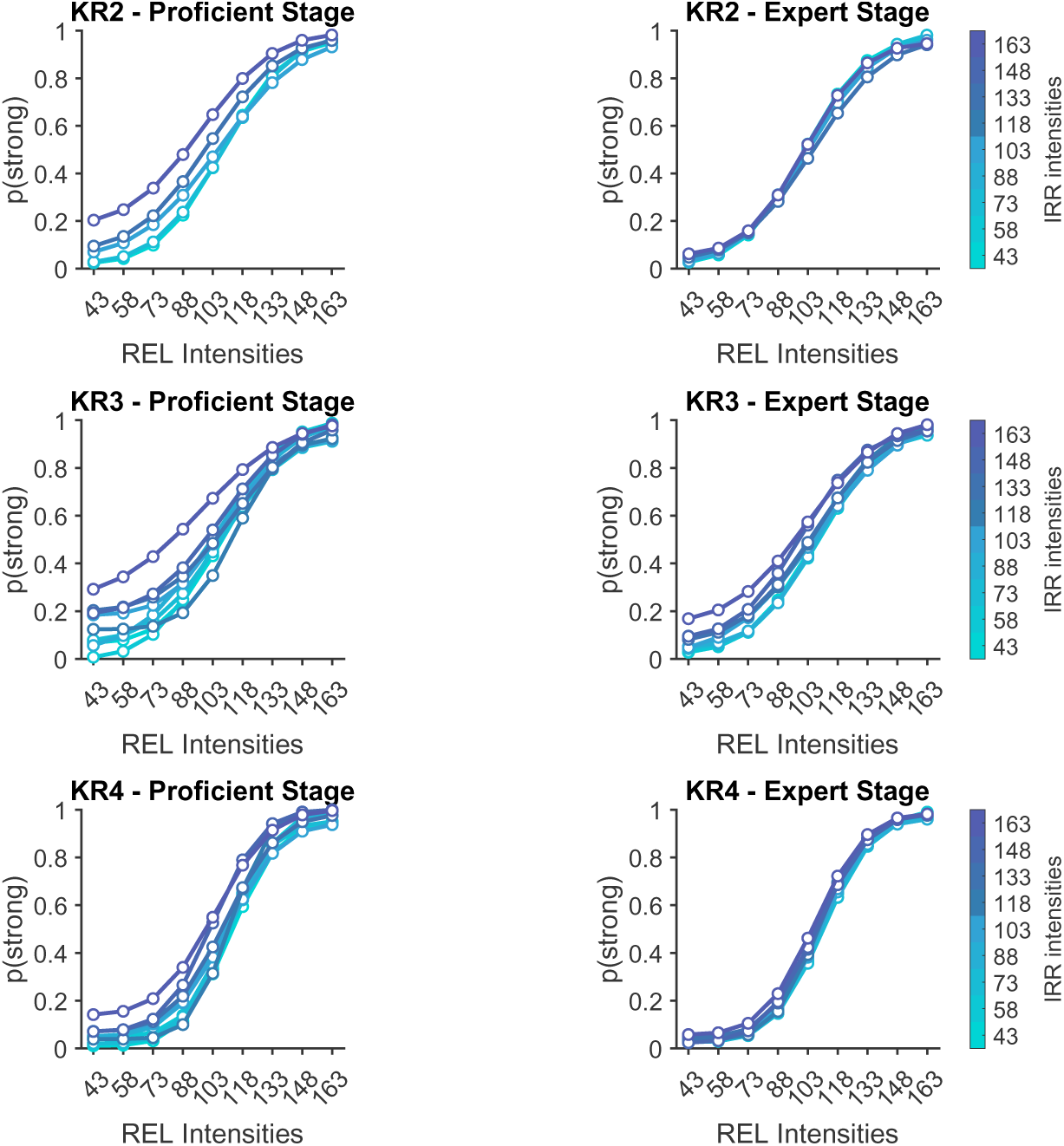
Psychometric functions identifying the proficient stage and expert stage. Difference in IRR bias during the two training stages, for rats KR2, KR3, KR4 performing the standard task.

**Figure 12:**
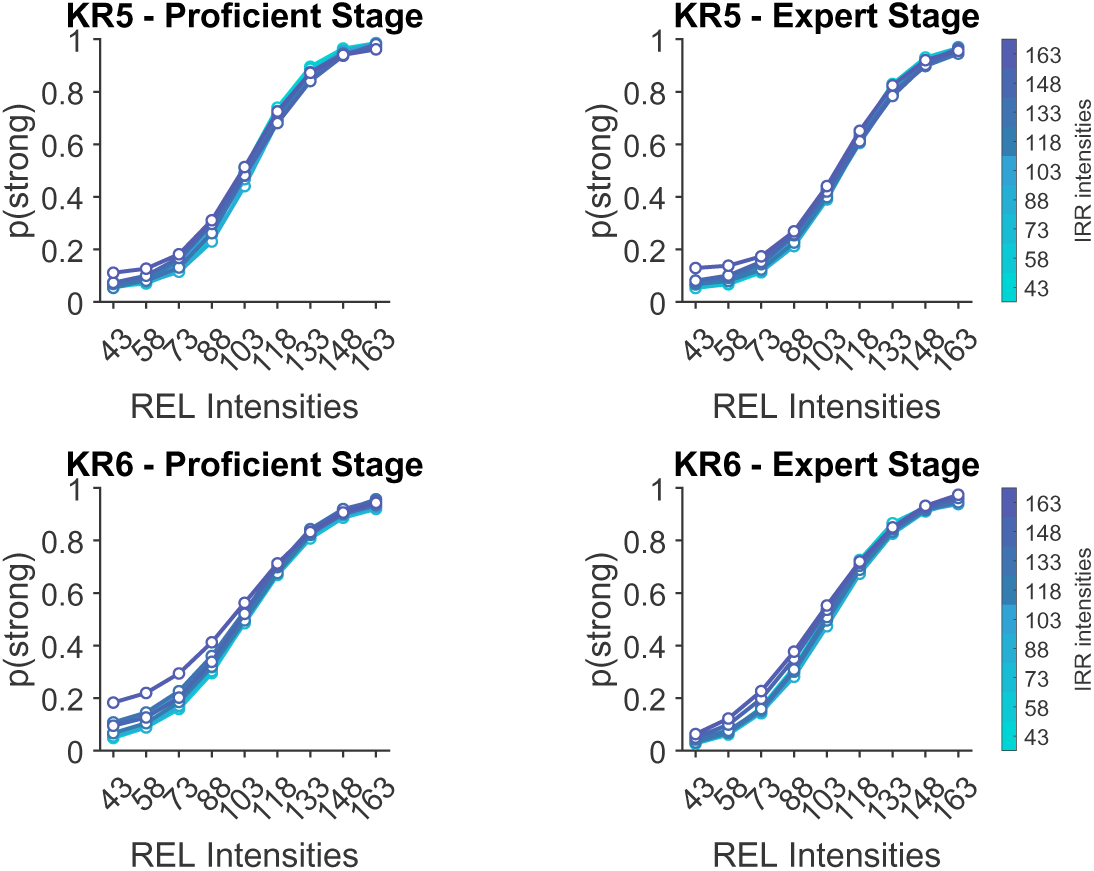
Psychometric functions identifying the proficient stage and expert stage. Difference in IRR bias during the two training stages, for rats KR5 and KR6 performing the standard task.

Video inspection during the experiment, and recorded video clips, showed that rats’ whiskers were in contact with the plate during both IRR and REL, signifying that the reduced weight of the IRR did not arise from preventing the vibration from being transduced by the sensory system.

Together, these results indicate that rats can learn to ignore irrelevant information when task structure is stable, but exhibit limited flexibility when relevance must be rapidly reassigned across time.

### Neural representations of relevant and irrelevant stimuli

#### Encoding of relevant and irrelevant stimuli in expert animals

To relate neural activity to behavioral performance, we recorded extracellular activity from motor (M1/M2) and somatosensory (vS1) cortices in four rats performing the task. These animals spanned different behavioral regimes: two expert rats (KR2, KR3), one proficient rat (MG1), and one rat trained in the dynamic task (KR8). This allowed us to relate neural dynamics to behavioral strategy and performance.

We first examined neural representations in expert rats performing the standard task. Behavioral analyses during recording sessions confirmed that these animals had effectively suppressed the influence of the irrelevant stimulus (Figure 13a-b).

**Figure 13:**
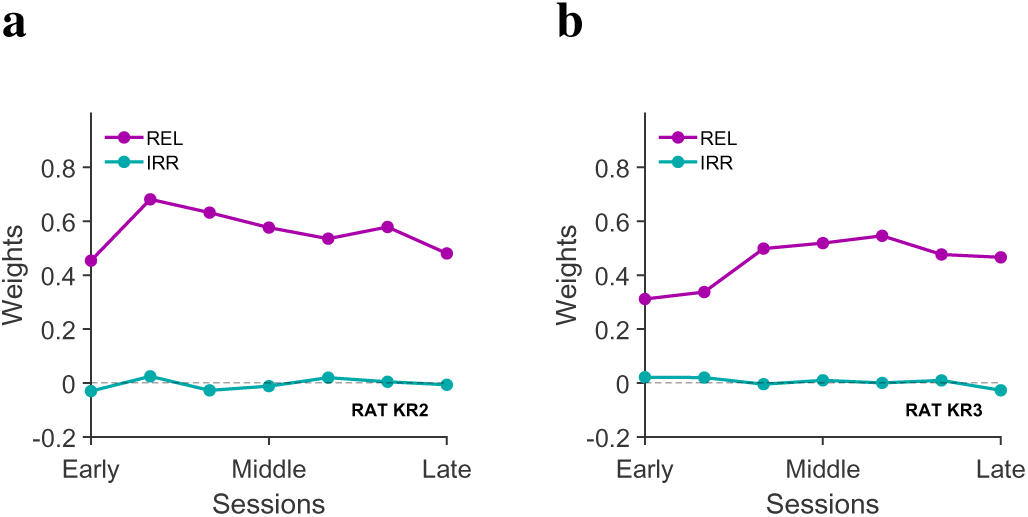
GLM weights during the recording sessions (standard task), for the two M1/M2-implanted expert rats. **a.** Weights of REL and IRR during the recording sessions, for rat KR2. **b.** Weights of REL and IRR during the recording sessions, for rat KR3.

Single-unit activity in M1/M2 showed modulation by both stimulus relevance (REL vs IRR) and intensity category (Figure 14), indicating sensitivity to task variables. At the population level, decoding analyses revealed that the intensity category of REL and the animal’s choice could be robustly predicted from M1/M2 activity in both rats (Figure 15). In contrast, the irrelevant stimulus (IRR) was either not decodable (KR2) or only weakly represented (KR3), particularly during the inter-stimulus interval.

**Figure 14:**
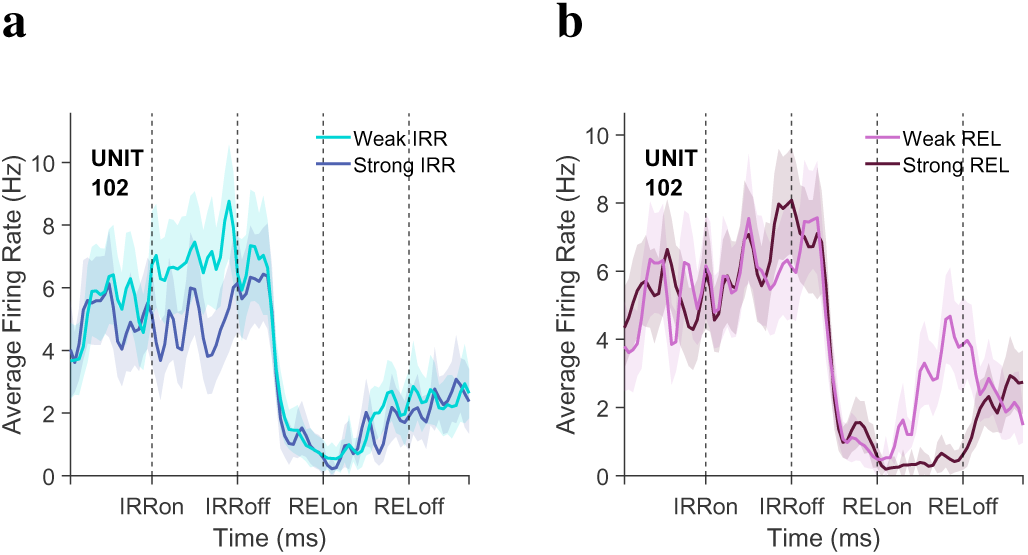
Example single unit’s firing rate. **a.** Average firing rate of example unit 102 during the entire trial, conditioned on IRR intensity category. **b.** Average firing rate conditioned on REL intensity category.

**Figure 15:**
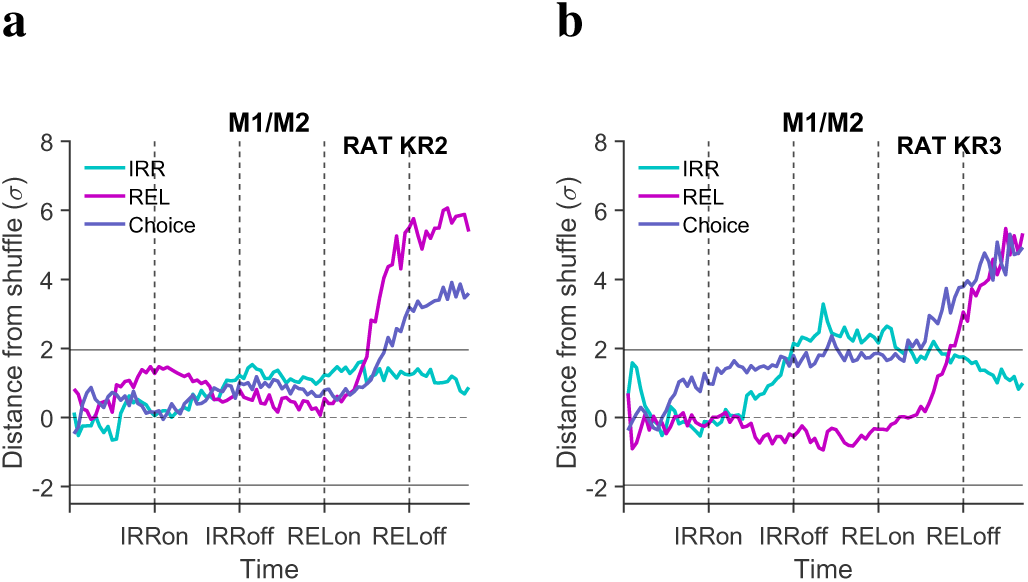
IRR, REL, and choice decoding in M1/M2, for rats KR2 and KR3. **a.** Decoding results for rat KR2. **b.** Decoding results for rat KR3.

#### Neural correlates of residual bias in a proficient animal

We next analyzed a proficient animal (MG1), characterized behaviorally by accurate judgment of REL in addition to a persistent IRR-induced bias (Figure 16). Recordings in both vS1 and M1/M2 revealed substantial differences from expert animals.

**Figure 16:**
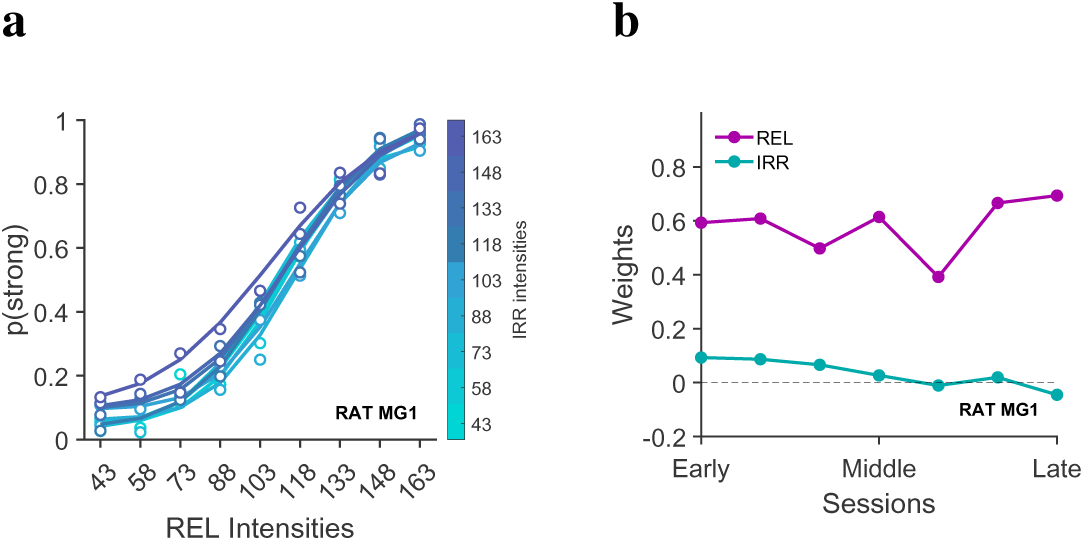
Behavioral performance of proficient rat MG1 during the recording period. **a.** Psychometric functions describing the IRR attractive bias. **b.** GLM weights of REL and IRR.

In vS1, both REL and IRR were significantly decodable (Figure 17a), indicating that irrelevant information was still encoded at the sensory level. In M1/M2, IRR remained decodable but showed partial suppression following REL onset, while REL was robustly represented (Figure 17b). Choice signals, however, were weak or absent compared to expert animals, suggesting incomplete transformation of sensory inputs into categorical decisions.

**Figure 17:**
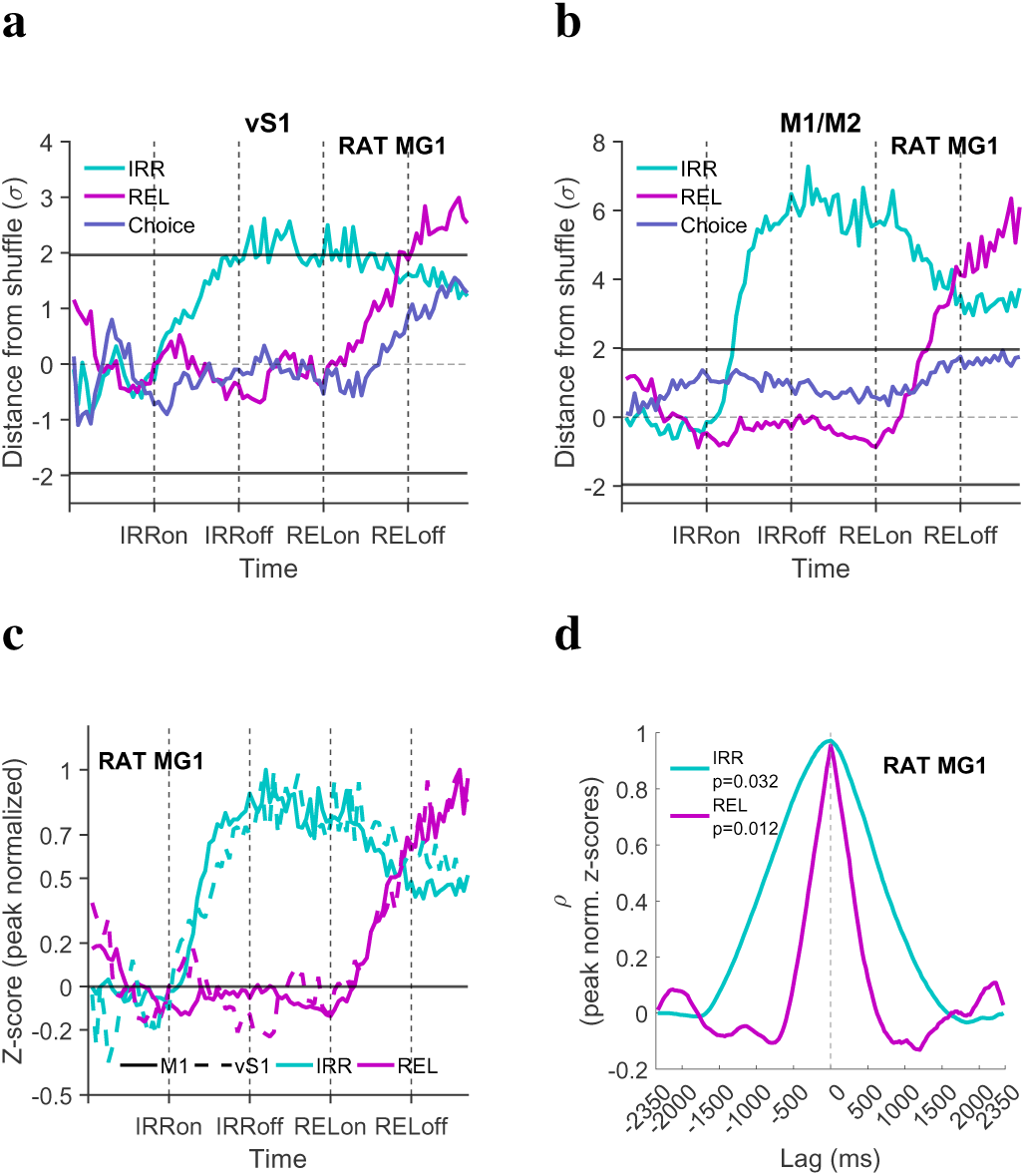
IRR, REL, and choice decoding in vS1 and M1/M2, for proficient rat MG1. **a.** Decoding in vS1. **b.** Decoding in M1/M2. **c.** Peak-normalized z-scores of IRR and REL decoding in both vS1 and M1. **d.** Lag analysis (cross-correlation of peak-normalized z-scores of vS1 and M1).

Comparing decoding magnitudes across areas revealed stronger signals in M1/M2 than in vS1, consistent with the emergence of more abstract representations in frontal cortex [11, 12]. Temporal analyses indicated that both areas encoded stimuli on similar time scales, with no consistent lag between them (Figure 17c-d), suggesting coordinated rather than strictly sequential processing.

#### Outcome-dependent modulation of stimulus representations

To probe how neural representations relate to behavioral success, we compared decoding during correct and error trials.

In expert animals, IRR decoding was minimal or transient during correct trials but could occasionally emerge during error trials (Figure 18). In contrast, REL remained decodable in both outcomes, but its representation differed qualitatively, suggesting that errors may reflect misrepresentation rather than absence of encoding.

**Figure 18:**
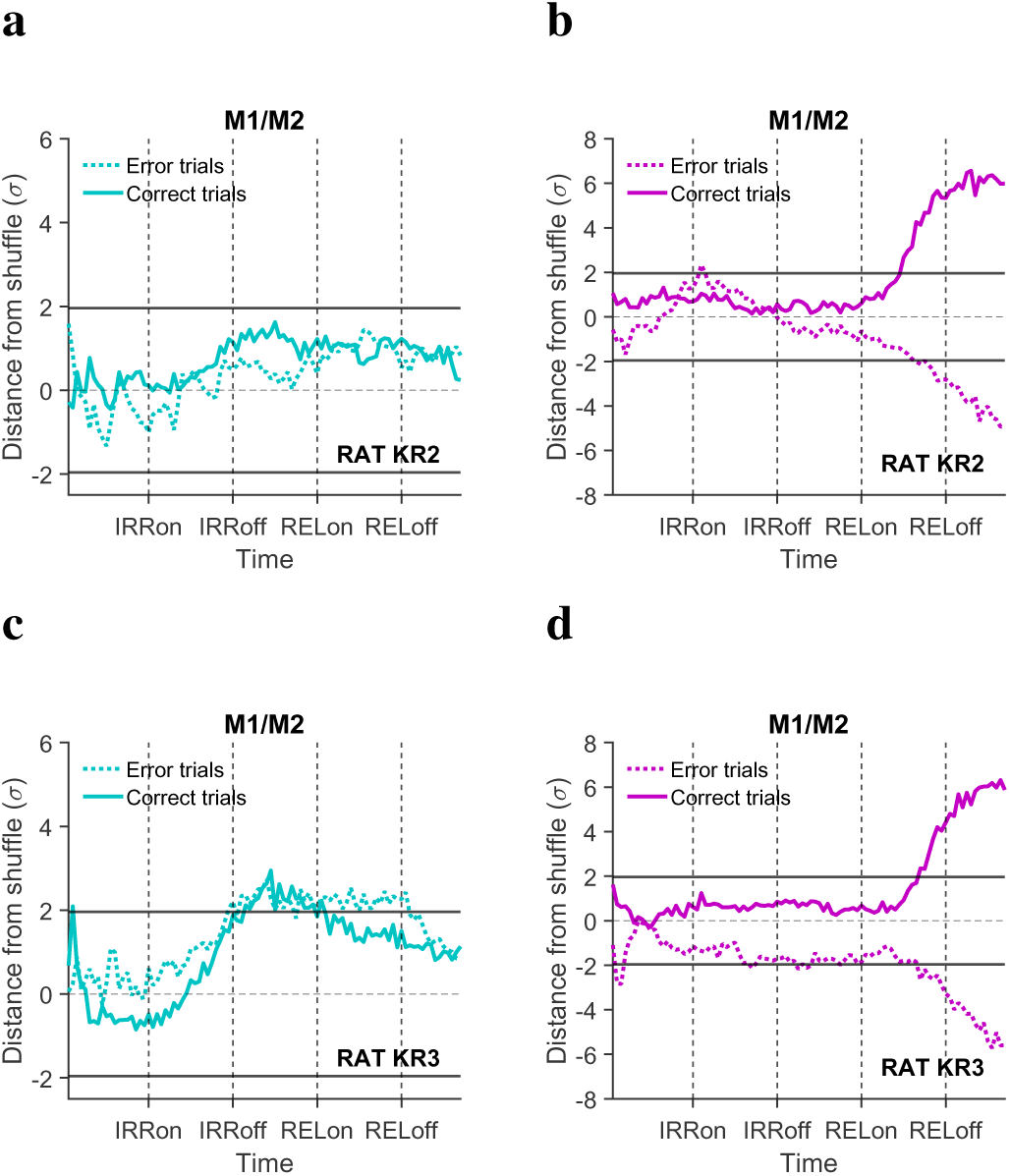
Decoding REL and IRR during correct and error trials, for rats KR2 and KR3. **a.** IRR decoding for rat KR2. **b.** REL decoding for rat KR2. **c.** IRR decoding for rat KR3. **d.** REL decoding for rat KR3.

In the proficient animal, outcome-dependent effects were more pronounced (Figure 19). In vS1, IRR was significantly decoded during error trials but suppressed during correct trials, consistent with attentional modulation at the sensory level. In M1/M2, both REL and IRR were generally represented, with stronger engagement during correct trials. However, during errors, both areas exhibited weaker and less consistent encoding of REL, indicating a broader failure of stimulus processing.

**Figure 19:**
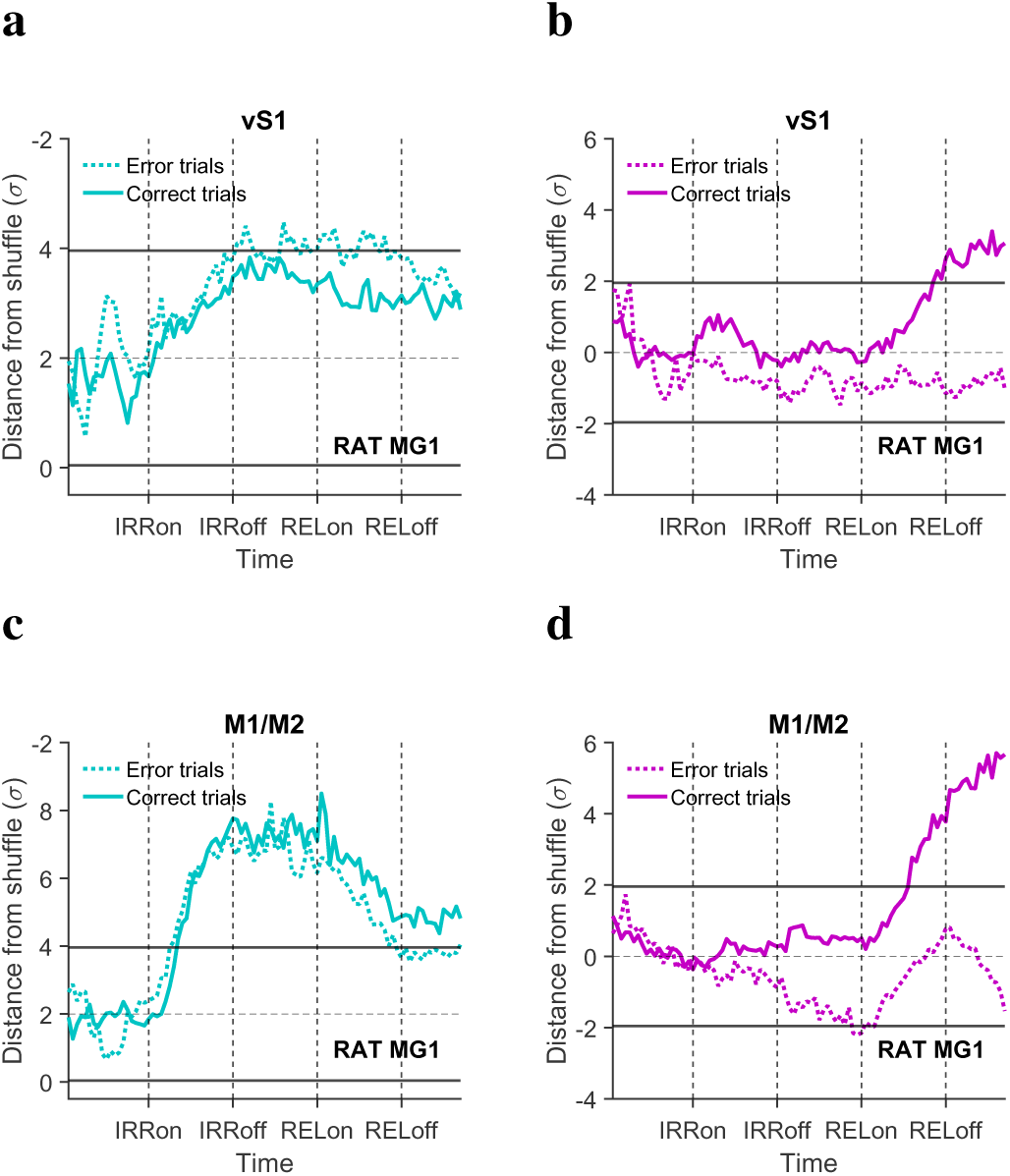
IRR and REL decoding in vS1 and M1/M2, during error and correct trials, for rat MG1. **a.** IRR decoding in vS1. **b.** REL decoding in vS1. **c.** IRR decoding in M1/M2. **d.** REL decoding in M1/M2.

Together, these results suggest that proficient performance is associated with partial suppression of irrelevant information and that errors reflect a broad disruption in stimulus representation and attentional modulation.

#### Misalignment between neural encoding and behavior in the dynamic task

Finally, we examined neural representations in a rat performing the dynamic task (KR8), which behaviorally relied on a suboptimal heuristic favoring the second stimulus (Figure 20).

**Figure 20:**
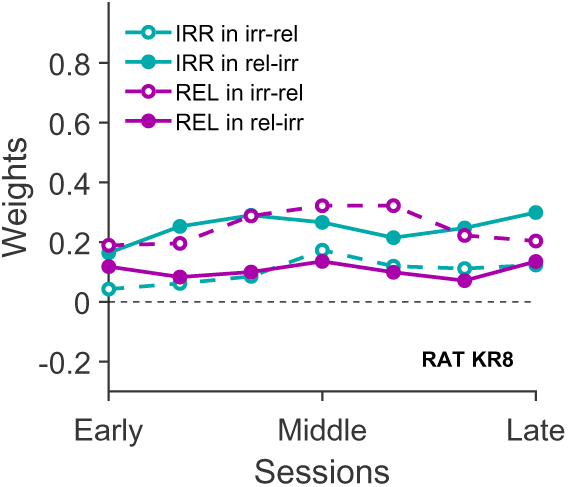
GLM weights during the recording sessions of rat KR8. GLM analysis of IRR and REL weights for REL-IRR trials and IRR-REL trials.

Decoding analyses revealed a striking dissociation between neural representations and behavior (Figure 21). In both vS1 and M1/M2, the first stimulus of the sequence—regardless of its relevance—was robustly encoded, whereas the second stimulus was only weakly or inconsistently represented. Choice signals were also weak and did not reach significance.

**Figure 21:**
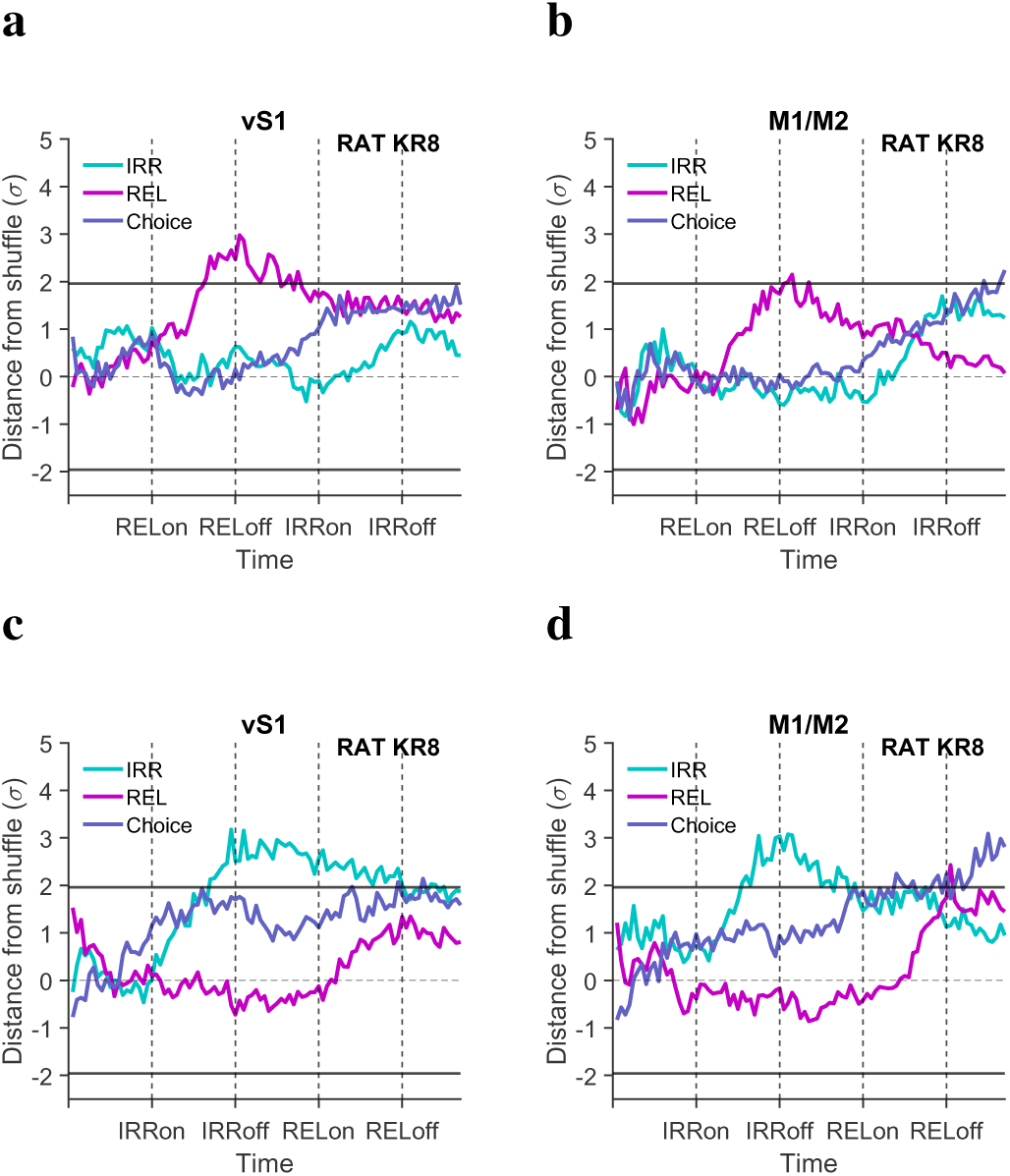
Decoding results in vS1 and M1/M2, for REL-IRR and IRR-REL trials, for control rat KR8. **a.** Decoding in vS1, for REL-IRR trials. **b.** Decoding in M1/M2, for REL-IRR trials. **c.** Decoding results in vS1, for IRR-REL trials. **d.** Decoding results in M1/M2, for IRR-REL trials.

This pattern indicates a misalignment between neural encoding and behavioral strategy: while neuronal populations preferentially represented the first stimulus, behavior was biased toward the second. These findings highlight that suboptimal behavioral strategies are associated with misaligned or weaker neural representations.

### Inter-areal interactions during task performance

So far, we have shown that neural representations of relevant and irrelevant stimuli differ across behavioral regimes. We next asked how interactions between sensory and frontal cortices contribute to this process, focusing on inter-areal synchronization and the direction of information flow.

To investigate how interactions between sensory and frontal cortices support task performance, we analyzed local field potentials (LFPs) in vS1 and M1/M2, focusing on synchronization and directional coupling.

#### Coherence reflects effective stimulus processing

In the proficient animal performing the standard task, vS1 and M1/M2 showed strong coherence across multiple frequency bands during stimulus processing (Figure 22b-d). In particular, gamma-band synchronization increased during stimulus epochs and was stronger for the relevant stimulus, consistent with a role in information transfer.

**Figure 22:**
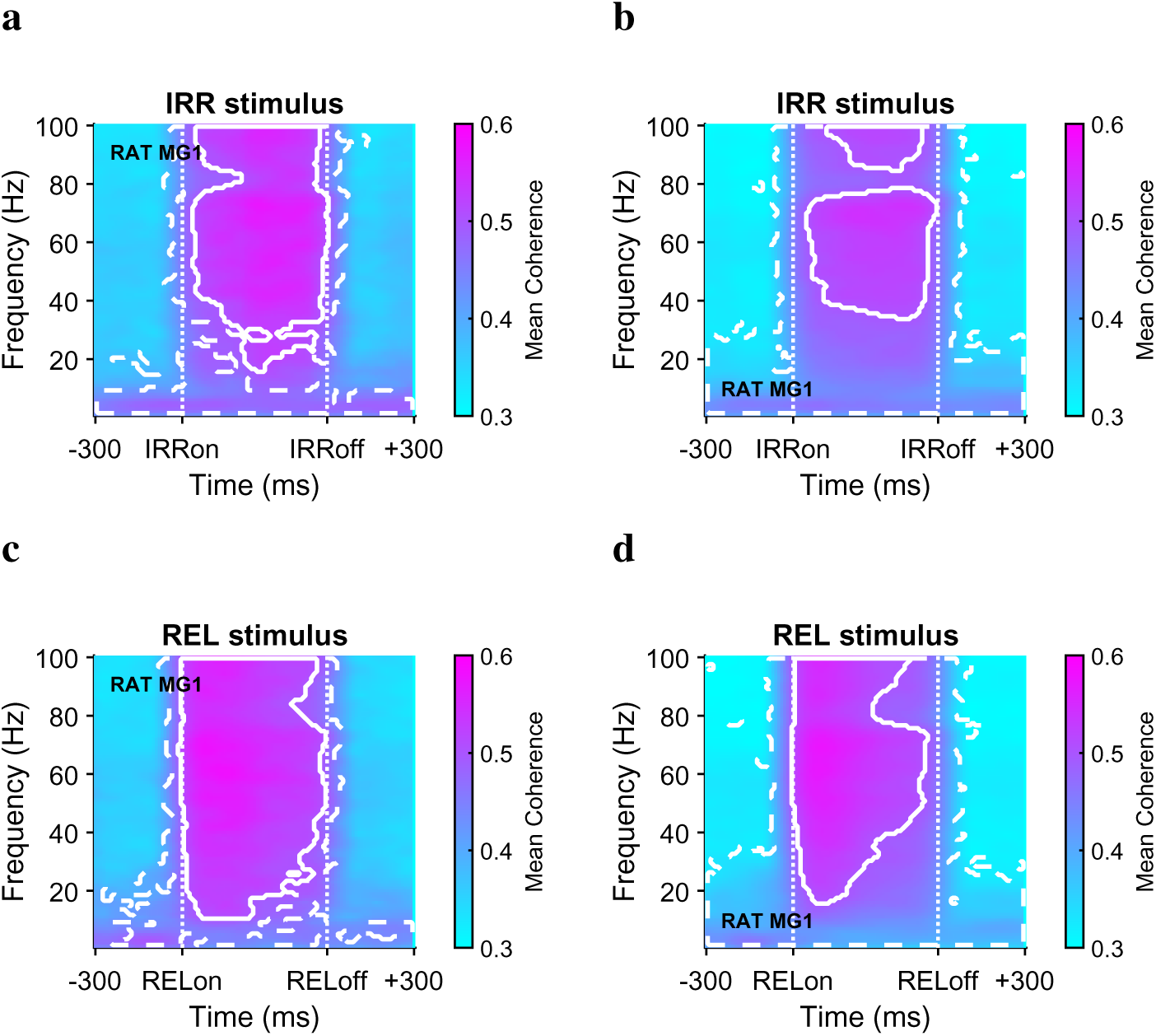
Coherograms during IRR and REL stimulus epochs, for incorrect and correct trials, for rat MG1. **a.** Coherogram for IRR stimulus epoch during incorrect trials (including 300 ms before stimulus onset, and 300 ms after stimulus offset). Dashed lines represent frequency ranges and time bins during which coherence was statistically significant. Solid lines represent frequency ranges and time bins during which coherence was higher than 0.5 **b.** Coherogram for IRR stimulus, during correct trials. **c.** Coherogram for REL stimulus, during incorrect trials. **d.** Coherogram for REL stimulus, during correct trials.

Interestingly, coherence increased further during error trials (Figure 22a-c), particularly in the gamma range. This hyper-synchronization may reflect inefficient or uncontrolled network interactions, indicating that stronger coupling is not necessarily beneficial for performance.

In contrast, in the dynamic-task animal, coherence between areas was markedly reduced (Figure 23). This reduction paralleled weaker decoding of both stimuli, supporting the idea that inter-areal synchronization contributes to effective stimulus representation.

**Figure 23:**
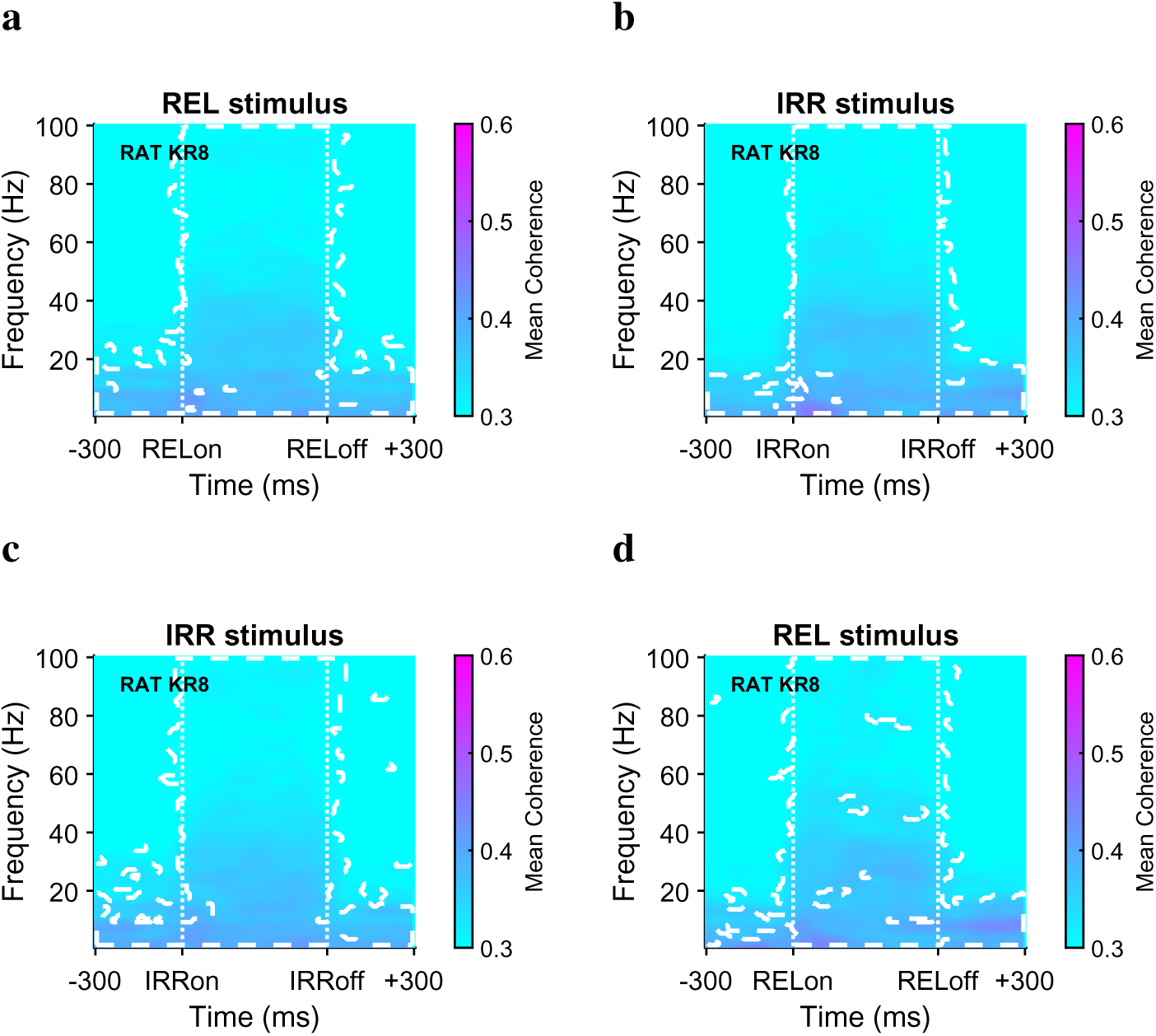
Coherograms during IRR and REL stimulus epochs, for REL-IRR and IRR-REL trials, for rat KR8. **a.** Coherogram for REL stimulus epoch during REL-IRR trials (including 300 ms before stimulus onset, and 300 ms after stimulus offset). Dashed lines represent frequency ranges and time bins during which coherence was statistically significant. **b.** Same, for IRR stimulus. **c.** Coherogram for IRR stimulus, during IRR-REL trials. **d.** Same, for REL stimulus.

#### Directionality of information flow: top-down modulation

We next examined the direction of information flow using Granger causality analysis. In the proficient animal, outcome-dependent analyses revealed than, during hits, both stimulus epochs were characterized by significant top-down modulation from M1/M2 to vS1 across a broad frequency range, particularly in beta and gamma bands (Figure 24b-d). This modulation during the stimulus presentation suggests that frontal cortex exerts control over sensory processing. By contrast, during error trials, top-down modulation was reduced (Figure 24a-c), especially in the beta band, indicating that effective top-down influence is associated with successful performance.

**Figure 24:**
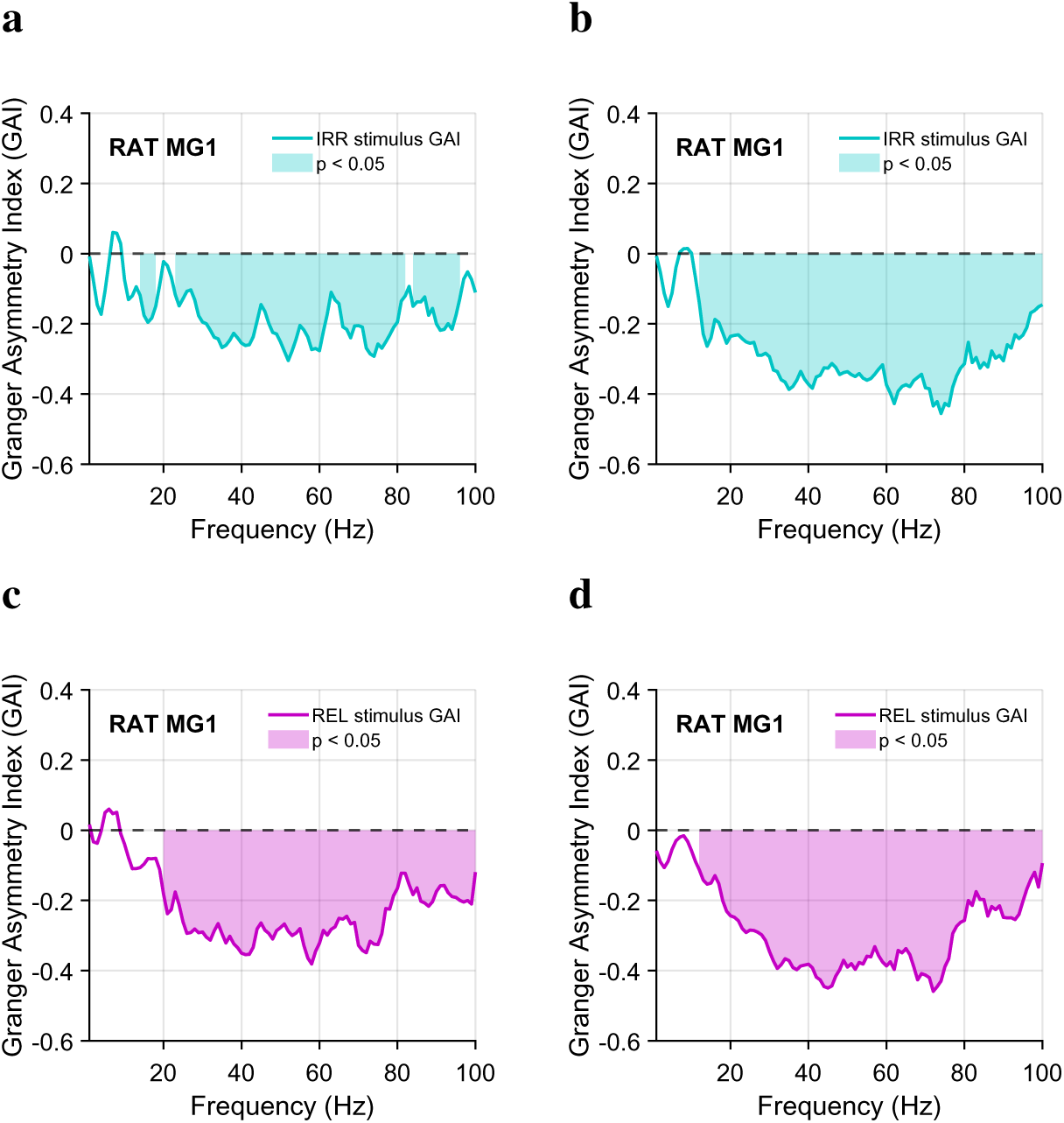
Granger Causality between vS1 and M1/M2, for both REL and IRR epochs, during misses and hits, for rat MG1. **a.** Frequency domain Granger Causality during IRR epoch, for error trials. Shaded areas represent frequencies in which GAI values were statistically significant (*p<0.05*, cluster-based permutation test). **b.** Same, for correct trials. **c.** Time-frequency domain Granger Causality during REL stimulus epoch, for incorrect trials. **d.** Same, for correct trials epoch.

In the dynamic-task animal, top-down modulation was present but altered (Figure 25). Beta-band influence emerged primarily during the second stimulus epoch, aligning with the animal’s behavioral bias toward the second stimulus. However, this modulation failed to support effective stimulus representation, as gamma synchronization remained weak.

**Figure 25:**
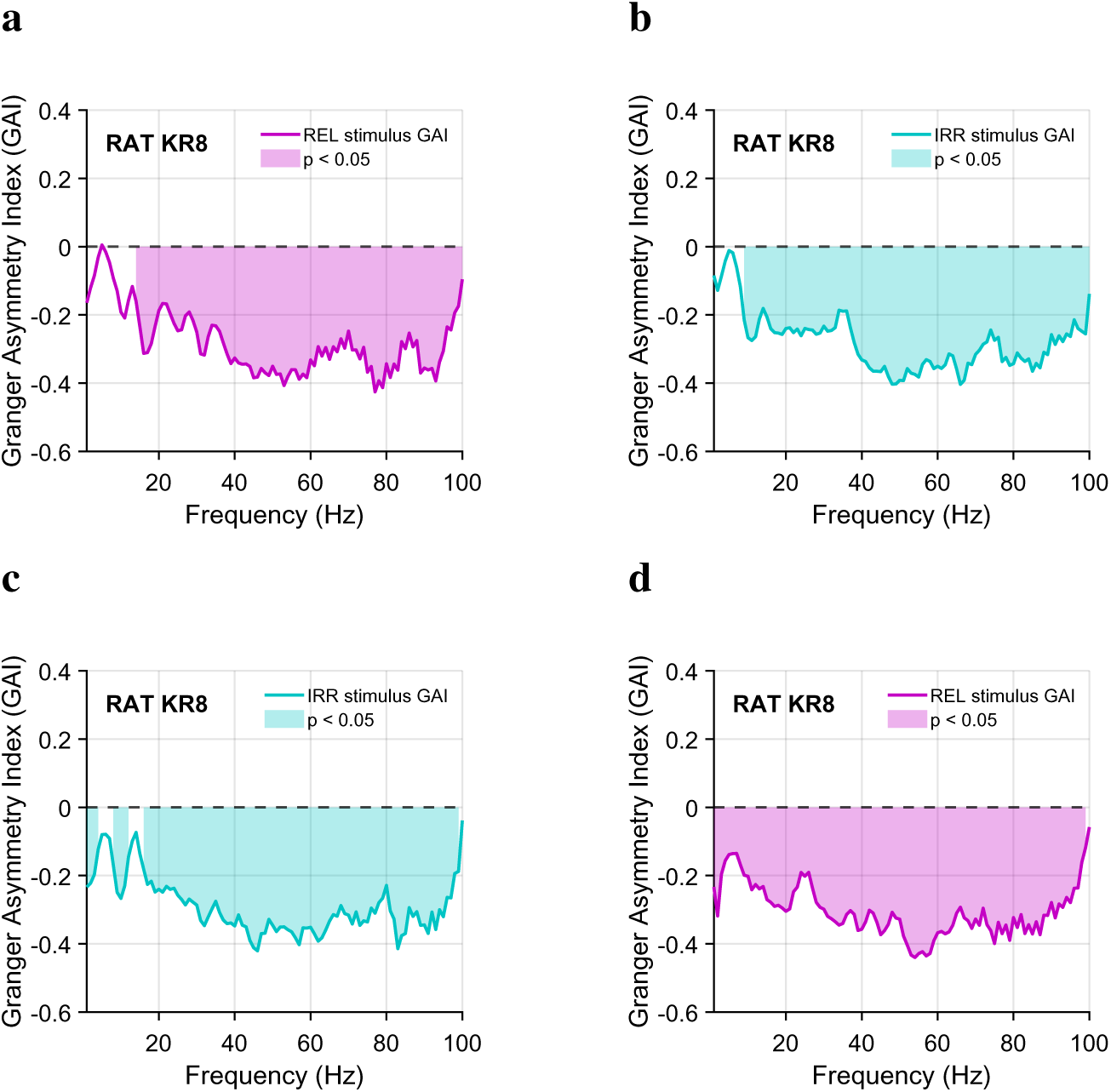
Granger Causality between vS1 and M1/M2, for both REL and IRR epochs, during REL-IRR trials, for rat KR8. **a.** Frequency domain Granger Causality during REL epoch, for REL-IRR trials. Shaded areas represent frequencies in which GAI values were statistically significant (*p<0.05*) cluster-based permutation test). **b.** Same, for IRR stimulus. **c.** Frequency domain Granger Causality during IRR epoch, for IRR-REL trials. **d.** Same, for REL stimulus.

#### Summary

Across animals, neural representations of relevant and irrelevant stimuli closely tracked behavioral strategies and levels of expertise. Expert performance was associated with strong encoding of relevant information and effective suppression of irrelevant inputs, whereas proficient performance reflected partial suppression and increased variability. In the dynamic task, suboptimal strategies were accompanied by a marked dissociation between neural encoding and behavior. At the circuit level, these differences were paralleled by changes in inter-areal synchronization and top-down modulation between frontal and sensory cortices.

## Discussion

### Irrelevant information biases perceptual judgments

In this study, we examined how rats allocate attention across time when required to ignore an irrelevant stimulus. Although animals successfully learned the task, their judgments were systematically biased by the irrelevant stimulus, which exerted an attractive effect on perception. This bias was strongest for weak relevant stimuli and decreased as a function of inter-stimulus interval (ISI), indicating that the interaction between stimuli is temporally constrained.

The asymmetry of the effect, with weaker relevant stimuli being more susceptible to interference, is consistent with differences in signal reliability. Weak stimuli generate less robust representations and are therefore more vulnerable to contamination by competing inputs [13,14].

### Learning reduces – but does not abolish – irrelevant information

Rats trained on the standard task progressively reduced the influence of the irrelevant stimulus. This is evident from the decrease in its contribution to behavioral choice over training. This reduction does not imply that irrelevant information disappears at the sensory level. Rather, the data suggest that its impact on behavior is attenuated.

Across animals, learning followed different trajectories. Some rats gradually reduced the influence of the irrelevant stimulus, while others showed little susceptibility from early in training. We therefore define two operational regimes: a proficient regime, in which behavior is accurate but still biased, and an expert regime, in which the bias is minimal.

In contrast, rats trained on the dynamic task did not achieve comparable suppression. When relevance varied across trials, animals did not rely on the acoustic cue but instead adopted a simpler rule, favoring the second stimulus. This suggests that while rats can learn to allocate attention across time under stable conditions, they exhibit limited flexibility when relevance must be reassigned dynamically.

This difference can be understood at the level of task demands. The standard task provides a stable mapping that supports learning of a consistent policy. The dynamic task introduces a conflict between temporal structure and task rule, increasing computational demands [15]. Under these conditions, animals appear to adopt a simpler temporal strategy that is only partially effective.

### Behavioral variability reflects constrained solutions

Despite variability across individuals, behavior was not random [16, 17]. Instead, it clustered into a small number of reproducible patterns. In the standard task, animals differed in the degree to which irrelevant information influenced their choices. In the dynamic task, all animals converged on a similar heuristic based on stimulus order.

These observations suggest that attentional control in this paradigm does not rely on a single solution, but instead emerges from a constrained set of strategies shaped by task structure [18–20]. Temporal order appears to act as a strong organizing principle, even when it is not explicitly informative.

The distinction between proficient and expert performance should be understood as an operational one. The data do not indicate discrete categories, but rather regimes within a continuum of behavior.

### Neural representations dissociate encoding from readout

To relate behavioral variability to neural dynamics, we recorded activity in M1/M2 and vS1 across animals expressing different behavioral regimes. Across these recordings, a consistent dissociation emerged between sensory encoding and behavioral readout.

In expert animals, M1/M2 robustly encoded the relevant stimulus and predicted choice, while the irrelevant stimulus was weakly represented or not decodable. However, the two expert rats differed in the extent to which irrelevant information remained detectable. One animal showed no clear representation of the irrelevant stimulus, whereas the other retained a residual encoding despite the absence of behavioral influence.

This dissociation suggests that suppression of irrelevant information operates primarily at the level of readout rather than through complete elimination of sensory encoding. In other words, irrelevant stimuli may still be represented neurally, but their influence on choice is reduced. Accordingly, similar behavioral outcomes can arise from different underlying neural states.

While these patterns are consistent with different computational solutions, such as selection-like versus integration-like processing, they do not uniquely specify the underlying mechanisms. Given the limited number of subjects and the indirect nature of decoding analyses, alternative explanations remain possible. In the proficient animal, both relevant and irrelevant stimuli were robustly encoded in vS1 and M1/M2. Partial suppression of the irrelevant stimulus emerged only after the onset of the relevant one, and choice-related signals were weaker and less stable than in expert animals. This suggests that proficient performance reflects a regime in which irrelevant information is still processed and only incompletely filtered.

Error analyses support this view. Failures were associated with both degraded representation of relevant information and inconsistent attenuation of irrelevant input, indicating that errors reflect a distributed breakdown of processing rather than a single deficit.

### Suboptimal strategies reveal dissociations between encoding and decision

In the dynamic-task animal, neural and behavioral patterns diverged in a striking way. Neural activity in both vS1 and M1/M2 preferentially encoded the first stimulus, whereas behavior was biased toward the second. This reveals a misalignment between early sensory processing and downstream decision mechanisms.

One possible interpretation is that sensory encoding is largely driven by stimulus order or salience, whereas decision processes apply an additional temporal weighting favoring more recent inputs. Under this account, the observed heuristic arises at the level of readout rather than encoding.

More generally, this finding suggests that suboptimal strategies are not simply weaker versions of optimal ones. Instead, they may reflect qualitatively different mappings between sensory representations and behavior.

### Inter-areal interactions support – but do not determine – performance

Efficient task performance was associated with increased coordination between vS1 and M1/M2. In the proficient animal, coherence between these regions increased during stimulus processing, particularly in the gamma band. However, synchronization was even stronger during error trials, indicating that more coupling is not necessarily beneficial [21, 22].

These results suggest a non-monotonic relationship between inter-areal coupling and performance, in which both insufficient and excessive synchronization can be suboptimal. In the dynamic-task animal, reduced coherence was accompanied by weaker stimulus encoding, consistent with the idea that coordinated activity supports effective processing. However, these observations remain correlational.

Analyses of directionality revealed top-down influences from M1/M2 to vS1, particularly during stimulus epochs. This influence was reduced during error trials, especially in the beta band, and may reflect engagement of task-related control processes [23, 24]. However, given the limited dataset and the nature of the analysis, this interpretation should be treated with caution.

Interestingly, in the dynamic-task animal, beta modulation was aligned with the second stimulus, mirroring the behavioral bias, yet did not improve representation. This suggests that patterns of inter-areal communication can reflect the adopted strategy, even when that strategy is suboptimal.

## Conclusions

Attentional control in this task emerges from the interaction between intrinsic sensory biases, learned behavioral strategies, and top-down modulation. Rats can reduce the influence of irrelevant information when task structure is stable, but show limited flexibility when relevance must change dynamically.

Crucially, similar behavioral performance can arise from different neural representations, and suboptimal strategies can involve mismatches between encoding and decision processes. These findings argue against a single canonical mechanism of attentional control and instead support a framework in which multiple constrained solutions coexist within a shared task structure.

### Limitations and future directions

Several limitations should be acknowledged. The number of subjects, particularly for electrophysiological recordings, is limited, and individual animals should be considered case studies rather than representatives of a broader population. The absence of vS1 recordings in expert animals further restricts interpretation across processing stages.

In addition, inferences about underlying strategies are indirect. Neural decoding reveals what information is represented, but not how it is used to guide behavior. As a result, different neural patterns may reflect multiple plausible computational mechanisms.

Future work should track learning within subjects, increase sample size, and incorporate causal manipulations to directly test the role of specific circuits and frequency bands in attentional control.

## Methods

All protocols were conformed to international norms and were approved by the Ethics Committee of SISSA and by the Italian Health Ministry.

### Rat subjects

9 Wistar male rats (Envigo) were housed in pairs and examined periodically by a veterinarian. They were regularly checked for their health and welfare conditions, and provided daily with environmental and social enrichment. They were maintained on a 12/12-hour light/dark cycle. The rats began the training protocols at the age of 8 weeks. To maintain motivation during the behavioral training, animals were water-deprived. The behavioral training protocol required approximately 1 hour of training on each week-day. During weekends, the rats had access to water for around 30 hours.

### Experimental apparatus

The experimental setup consisted of a T-shaped Plexiglas box, measuring 25 x 25 x 38 cm. In the front wall, a hole allowed the rat to extend its head into the stimulus delivery port. Here, a nose poke detected the snout by an infrared detector, while a blue LED light signaled to the rat that a new trial could begin. The setup was positioned into a Faraday cage, and the experiments were run the in the dark. The only light provided was a red LED, and was used by the experimenter to monitor the rats during the task, via a webcam (Logitech HD Webcam C310).

Two reward spouts, on 8 cm high pedestals, were placed to the right and left compared to the nose poke. Two custom made infrared sensors, one for each spout, detected the rat’s position and activated reward delivery accordingly. Water was delivered by two syringe-pumps, one for side, controlled by a AVR32 board (National Instruments, Austin, TX).

Three audio speakers were placed on the three walls of the setup, one above each spout and one at the back. The one at the back delivered the go cue, while the lateral ones gave a reward delivery cue, upon correct response, to act as reinforcement. Two additional audio speakers were placed on the front external sides of the setup: these delivered the acoustic cue used to tag the relevant vibrotactile stimulus. Experiments were controlled using in-house LabVIEW software (National Instruments).

### Vibrotactile stimulus generation

The stimulation medium consisted of a rectangular plate measuring 20 x 30 mm connected to a motorized shaker (Bruel and Kjar, type 4808), to which velocity values were sent as analog signals, moving the plate along the rostro-caudal axis. Stimuli were vibrations of the plate made of low-pass filtered Gaussian white noise, obtained by stringing together randomly sampled velocities. Velocities were obtained by sampling a normal distribution with 0 mean and defined by the standard deviation *σ*, ranging from 2.84 to 14.85. There were 50 seeds available for each velocity standard deviation. The noise was low-passed through a Butterworth filter with 150 Hz cutoff, amplified, and sent as voltage input to the shaker motor. It is possible to convert the intensities values defined by *σ* in mean speed values (sp, measured in mm/s) by using the following equation:

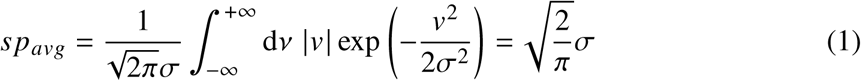

In every session, there were 9 or 5 linearly-spaced values of sp.

### Experimental design: Irrelevant Stimulus Paradigm

Before every trial, the blue light turned on to signal to the rat that a new trial could begin. The rat had to insert its head in the hole and keep its nose in the nose poke, triggering the delivery of stimuli. The rat had to remain in the nose poke for the entire trial, before a go cue signaled the rat to make a choice, by exiting the nose poke and turning to the left or right sides of the setup, based on the rule that was reinforced during learning. Half of the rats, randomly chosen, were trained to go to the left spout if the relevant stimulus was stronger than the boundary value, the other half were trained following the reversed rule.

The task was designed in the following way: pre-stimulus delay (500 ms), irrelevant stimulus IRR (500 ms), inter-stimulus interval (ranging from 500 ms to 2000 ms), relevant stimulus REL (500 ms), post-stimulus delay (500 ms). The acoustic cue tagging the relevant stimulus lasted 300 ms and overlapped the last 300 ms of the ISI. In some instances (e.g., early withdrawals from the nose poke in highly trained rats), the post-stimulus delay was reduced to 350 ms, to reduce the number of early withdrawals and the number of trials excluded from the analysis. This also helped the animals to be less nervous and calmer during the behavioral training.

9 stimulus intensities were employed for both the relevant and irrelevant stimuli, and could be combined in 81 possible different permutations. Only one rat (KR2) had a different stimulus set (9 intensities for REL, 5 intensities for IRR), as it was initially tested as a pilot subject. Behavioral and neuronal results of this rat were consistent with the other subjects, excluding the possibility that a different stimulus set led to different results.

#### Standard task

The standard task represented the first implementation of the Irrelevant Stimulus Paradigm above described. In the standard task, the first stimulus of the sequence is IRR, and the second is REL.

Rats KR2, KR3, KR4, KR5, and KR6 were trained performing the standard task.

#### Control task

In this task variant the acoustic cue tagging the relevant stimulus was removed. Rats KR2, KR3, KR4, and KR6 performed the control task.

#### Dynamic task

The dynamic task is analogous to the standard task, however, approximately half of the trials within a single training session were characterized by REL in the second temporal position, and the remaining approximate half of trials had REL in the first temporal position. The order of presentation of such trial types (IRR-REL and REL-IRR) was random.

Rats KR7 and KR8 were exclusively trained in the dynamic task. Rats MG1 and MG2 were initially trained in the dynamic task, and then trained exclusively with the standard task. Rat MG1 changed training regime approximately 1.5 months before performing neuronal recordings. This ensured that the rat achieved a proficient level of performance in the standard task.

### Behavioral analysis

#### Psychometric curve estimation

All analyses were performed in MATLAB (MathWorks, Natick MA) using custom code. To estimate psychometric functions, we computed for each stimulus vibration *sp*, the proportion of trials in which the rat responded by going to the side corresponding to the “strong” category. We fitted the data with a cumulative Gaussian logistic function, including asymmetric lapse parameters:

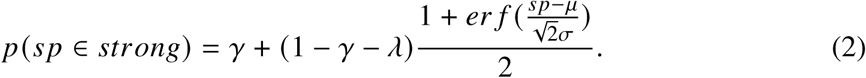

where *sp* is the stimulus intensity, *μ* is the midpoint parameter, *σ* is the slope parameter, *γ* and *λ* the lower and higher asymptotes of the function, respectively, corresponding to lapse rates. Parameters values were estimated by MLE using the MATLAB functions fmincon and fminsearch.

#### Generalized Linear Model estimation

GLM analyses were implemented using custom-made MATLAB code, fitting free parameters with the following equation:

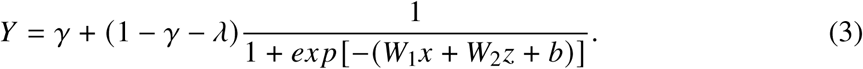

Where *Y* is the predicted rat’s choice, *γ* and *λ* are, respectively, the lower and upper lapse rates, *W*_1_ is the weight for the REL stimulus, *W*_2_ is the weight for the IRR stimulus, *x* is the REL stimulus intensity, and *z* is the IRR stimulus intensity. Parameters values were optimized by MLE using the MATLAB functions fmincon and fminsearch.

### Surgery protocol

For implanting electrodes into left M1/M2 and left vS1, both contralateral to the right side vibrissal stimuluation (M1/M2 only rats n=2, M1/M2 and vS1 rats n=2), animals were anesthetized with Isoflurane through a plastic snout mask, and was administered via a veterinary apparatus (V-1 Tabletop, VetEquip Inc.). The concentration of the agent was variable, based on the depth of required anesthesia and subject differences. To induce anesthesia, the rat was placed in a box and gradually anesthetized with Isoflurane until reaching an agent level of 2%-2.5% mixed with oxygen (O2 98%), as calculated according to the MAC value (Minimum Alveolar Concentration: 1.35% ± 0.10%). Once immobile, a mask was positioned on the rat’s snout to provide a constant administration of Isoflurane with concentration of 2.5%, and a flow rate of 1 l/min. This established a state characterized by analgesia, muscle relaxation, and loss of reflexes (absence to reaction to tail, hind paw pinches, and corneal air puff). This state was maintained across the entire surgical procedure (opening of skin and bone). Blood oxygen saturation was at least 98% and cardiac frequency was 350 ± 25 BPM.

Once the craniotomy/craniotomies was/were completed, provided that the rat continued to show no reflexes or signs of discomfort, Isoflurane concentration was decreased to 2% in 1 l/min of oxygen to prevent gas accumulation and allow a more rapid postoperative recovery. This same flow rate was maintained during the electrophysiological recordings used to verify electrode positions. At the end of the surgical procedure the Isoflurane vaporizer was turned off and the rat was allowed to recover while Isoflurane 1 l/min of oxygen. During the entire procedure and until the rat completely recovered from the anesthesia, body temperature, blood oxygenation, breathing frequency, and heart rate were continuously monitored. The absence of reflexes was regularly tested. In case of bleeding, fluids were reintroduced by subcutaneous injection to ensure proper hydration. Target cortical regions were accessed by craniotomies, using the Paxinos Rat Brain Atlas’ stereotaxic coordinates [25], and standard stereotaxic techniques. The dura mater was removed over the entire craniotomy with a small needle.

The electrode arrays were inserted by slowly advancing a Narishige automatic micromanipulator. After inserting the array, the remained exposed cortex was covered with biocompatible silicone (KwikSil; World Precision Instruments). In all rats, at least 4 small screws were fixed in the skull as support for dental cement. All screws served as ground and reference electrodes.

Before the conclusion of the surgical procedure, animals were given antibiotic (Baytril; 5 mg/kg; i.p) and analgesic (Rimadyl; 2.5 mg/kg; i.m.). In addition, both the antibiotic and the analgesic were delivered through a water bottle for 24 hours after the completion of the surgery. During this recovery time, rats had *ad libitum* access to water and food.

For the M1/M2-vS1 implanted rats, we decided not to proceed with the mapping of vS1 using a single probe, for the following reasons: i) extended surgical times pose a health risk to the animal, and should be kept as short as possible, ii) the microdrive array was designed in such a way to cover the greatest cortical area possible, to maximize the probability of placing individual electrodes in individual barrels, iii) the task requires the rat to be freely moving and not head fixed, making it not possible determine with certainty whether the stimulated whiskers were those always in contact with the stimulation plate.

#### Microdrives configuration

Rat KR2 was implanted with a microdrive array constituted by 16 electrodes, arranged in 2 rows of 8 electrodes, with the long side placed along the medio-lateral axis, targeting M1/M2. The stereotaxic coordinates of the craniotomy were 1 mm to 5.5 mm AP from bregma, and 0.5 mm to 4.25 mm ML from bregma.

Rat KR3 was implanted with a microdrive array constituted by 32 electrodes, arranged in 4 rows of 8 electrodes each, with the long side placed along the antero.posterior axis, targeting M1/M2. The stereotaxic coordinates of the craniotomy were 0.5 mm to 3.5 mm AP from bregma, and 0.5 mm to 3.5 mm ML from bregma.

Rats KR8 and MG1 were both implanted with two custom-made microdrive arrays constituted by 12 electrodes each, arranged in 3 rows of 4 electrodes. Both arrays had movable electrodes, meaning it was possible to move them deeper or shallower during the electrophysiological recordings period, to improve recording quality. The array targeting M1/M2 was placed having the long side aligned to the antero-posterior axis, whereas the vS1 array had the long side aligned to the medio-lateral axis. The stereotaxic coordinates for the craniotomy in M1/M2 were 0.5 mm to 2.5 mm AP from bregma, and 0.5 mm to 2 mm ML from bregma. For the vS1 craniotomy, the stereotaxic coordinates we used were -1 mm to -3.5 mm AP from bregma, and 4 mm to 7 mm ML from bregma.

### Neuronal analysis

#### Neuronal recordings preprocessing

Neuronal recordings were spike-sorted manually using Wave Clus [26]. Single units were selected according to spike shape consistency and inter-spike interval. To be included in the analyses, units had to meet two criteria: i) at least 2000 total spikes, and ii) firing in at least two trials per REL-IRR intensities combinations.

#### Firing rate

Firing rates were computed by binning spikes in non-overlapping 25 ms time-bins. A Gaussian smoothing over a 500 ms window with a standard deviation of 100 ms was then applied. Since trials had variable ISI duration, we only included the first 200 ms and last 300 ms of the ISI, overlapping with the delivery of the acoustic cue. Therefore, trials included pre-stimulus delay, IRR/REL, first 200 ms of ISI, last 300 ms of ISI, REL/IRR, post-stimulus delay.

#### Decoding

##### Pseudopopulation construction

The decoding analysis was performed using a pseudopopulation approach, since units were recorded across multiple sessions. We ran separate decoders for stimuli and choice. When decoding REL and IRR, for each recording session, trials were categorized into four conditions based on the factorial combinations of REL and IRR intensity categories. When decoding choice, trials were instead categorized according to the factorial combinations of REL intensity category and choice category. After labeling trials, we randomly split them in a training set and a test set (80% and 20%, for each condition). To account for unequal trial counts, and to diminish potential noise correlations, we used a bootstrapping approach, resampling with repetition 800 trials from the training set, and 200 trials from the test set. This procedure was performed independently for each unit and each session. Each resulting pseudosession of *N* units was concatenated to build a pseudopopulation where each condition was perfectly balanced. We started by including all trials (correct and incorrect trials), before running separate decoders for miss and hit trials.

##### Classifier

Before training the Linear Discriminant Analysis classifier (LDA), we projected the neuronal activity of the training set and test set onto their stimulus (or choice) first principal component, separately computed using dPCA [27]. LDA identifies a weight vector that maximizes the separation between condition means, while minimizing within-condition variance: we trained the classifier on the projected neural activity belonging to the training set, and then tested the model on the unseen test activity. Next, we obtained raw accuracy scores, and we assessed the classifier’s performance by computing the decoder’s score

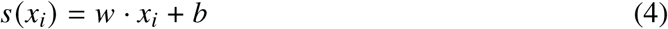

where *w* is the weight vector, *x_i_* is the vector of neural activity for trial *i*, and *b* is the bias term. The final classification for each test trial was determined by the sign of the score *s*(*x_i_*).

A separate LDA was trained and tested on projected neuronal activity with shuffled trial labels, to obtain a control distribution and shuffled accuracy scores.

This whole procedure was iterated 100 times, each time ensuring a different train/test split.

##### Statistical significance of decoding results

To test whether accuracy scores were statistically significant (i.e., statistically different from shuffled accuracy), we z-scored the real accuracies against the shuffled accuracies, similarly to [28]:

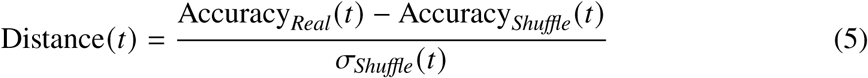

where Distance(*t*) is the z-score accuracy score for time *t*, Accuracy*_Real_*is the average accuracy over all iterations of cross-validation for correctly ordered labels, Accuracy*_Shuffle_* is the average accuracy over iterations for shuffled labels, and *σ_Shuffle_* is the standard deviation over all iterations for shuffled labels.

##### Relating classifier’s score and rat choices

For each iteration of the decoder, we stored the rat’s choice (i.e., answering “strong” or “weak”) to REL and IRR for each held-out trial, along with the classifier’s score, computed as described above. Next, we computed *p*(*strong*) for REL and IRR as proportion of “strong” choices, obtaining one probability value for each of the 100 iterations of the decoder. We averaged the single-trial classifier’s scores, for each iteration. Next, we chose the time-bin with the highest accuracy score for REL and IRR intensity category prediction, and plotted the averaged classifier’s score for strong/weak REL and IRR, against rat’s choices (*p*(*strong*) to REL and IRR).

##### Classifier for correct and incorrect trials

To visualize differences in stimulus processing related to correct or incorrect trials (i.e., trials in which the rat’s choice was rewarded or not), we performed a separate decoding analysis. The only difference lies on the trial pool used to train and test the classifier. When decoding hit trials, we pooled together only the hit trials, and then split into training/test. This ensured that the resulting accuracies reflected exclusively neuronal activity underlying optimal behavioral performance. When decoding miss trials, we built the training set by randomly sampling 80% of the hit trials. For the test set, we used the entire pool of miss trials. Given that error trials constitute approximately 25% of the whole trial set, we decided to use all error trials for the test set, in order to avoid underfitting.

##### Cross-correlation

We cross-correlated the vS1 and M1/M2 peak-normalized decoding z-scores using the MATLAB xcorr function to identify the optimal lag and the maximum correlation coefficient (*R_obs_*). Statistical significance was assessed using a permutation test (*n* = 1000 iterations). The null distribution was constructed by circularly shifting the vS1 z-scores while keeping the M1/M2 sequence fixed. This preserved the signal’s intrinsic autocorrelation structure while destroying the temporal alignment between areas. The p-value was defined as the proportion of permutations where *R_null_* ≥ *R_obs_*.

#### Local Field Potential acquisition and analysis

We performed inter-areal synchronization analysis and Granger causality analysis using the Fieldtrip toolbox [29] and custom-built MATLAB codes.

##### LFP acquisition and preprocessing

We obtained LFPs offline using neuronal data yielded by the same electrodes used for the decoding analysis, to ensure consistency and sufficient quality of recordings. Raw signals were downsampled to 1000 Hz, and bandpass filtered between 1 and 300 Hz. Since recordings were carried out within a Faraday cage (see Experimental apparatus), we abolished significant power-line interference, and therefore, chose not to apply a notch filter, preserving the integrity of the signal within the 50-60 Hz frequency range. We divided continuously recorded data in stimulus epochs corresponding to REL and IRR delivery, storing the timestamps of nose poke entry and exit. Since the ISI was variable, we could not segment the data in trials of identical duration, hence the stimulus epoch segmentation strategy. By storing the timestamps of nose poke entry/exit, we were able to include or exclude for further analyses activity occurring before or after the stimulus delivery.

##### Coherograms

To visualize the synchronized activity of vS1 and M1/M2, we computed time-frequency resolved coherograms for REL and IRR stimulus epochs, including 300 ms before stimulus onset, and 300 ms after stimulus offset. We used a Hanning taper to estimate power and cross-spectral density for frequencies ranging from 1 to 100 Hz, in 1 Hz steps. For all frequencies, we used a fixed 300 ms sliding window, which shifted in 10 ms increments across the considered epoch. We employed the 300 ms sliding window as it provided an optimal compromise between temporal and frequency resolution across the entire frequency range. It allowed the capturing of multiple cycles of low-frequency oscillations (*θ* and *β*), while maintaining sufficient temporal precision to track the dynamics of high-frequency ranges (*γ*), minimizing spectral leakage. We first computed the coherence for every possible pair-wise combination of electrodes between the two areas, and then averaged these individual channel-pair coherograms to produce a grand-averaged coherogram, providing a robust measure of the global synchronization between vS1 and M1/M2.

##### Frequency-resolved Granger causality

To identify the direction of information flow between vS1 and M1/M2, we ran frequency-resolved Granger causality. We included 500 ms from stimulus onset until stimulus offset (to include only stimulus epoch), for both REL and IRR. We achieved spectral decomposition using the Multitaper Fast Fourier Transform (mtmfft) with a Hanning taper, including frequencies ranging from 1 Hz to 100 Hz, in 1 Hz steps. To achieve a consistent frequency resolution of 0.25 Hz, we zero-padded all data segments to a 4 s duration prior to the FFT. We then estimated Granger causality for both directions (vS1 → M1/M2 and M1/M2 → vS1). To quantify the direction of information flow we used the Granger Asymmetry Index (GAI), which is the normalized difference between Granger Causality scores (GC) for each direction:

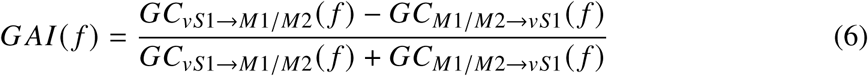

##### Time-frequency resolved Granger causality

We computed time-frequency resolved Granger causality using a sliding window with a duration of 300 ms, shifted in 10 ms increments across the epoch (from -300 ms to +800 ms relative to stimulus onset). Data segments were extracted with an additional 150 ms of padding on either side of each window, to ensure that all time-frequency estimates within the analyzed period were based on complete and non-truncated activity windows. We performed the spectral decomposition for each temporal window, by estimating Fourier coefficients using the Multitaper Fast Fourier Transform (mtmfft) with a Hanning taper, including frequencies between 1 Hz and 100 Hz, in 1 Hz steps. Each 300 ms segment was zero padded to 4 seconds prior to the FFT, to ensure a consistent frequency resolution of 0.25 Hz. We then computed the GAI:

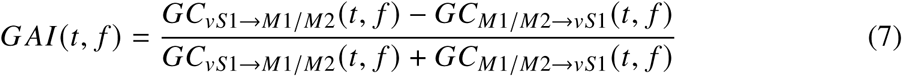

##### Cluster-based permutation tests

We used a non-parametric cluster-based permutation test to assess statistical significance when computing coherograms. The observed coherence values were compared against a null distribution generated by shuffling trial identities. Initial clusters were formed by grouping adjacent time-frequency bins exceeding a t-threshold corresponding to a *p* < 0.001 significance level. The significance of these clusters was then evaluated by comparing their summated t-values against a distribution created from 5000 random permutation. The final significance for a cluster was set at *p* < 0.05. This method controls for multiple comparisons.

For the frequency-resolved Granger causality, we used a similar approach. To assess whether the directionality was statistically significant against a null hypothesis of asymmetry, we used a non-parametric cluster-based permutation test across sessions, using a dependent-samples t-statistic. For each session, the observed GAI was compared to a null distribution of zeros. Initial clusters were formed by grouping adjacent frequency bins with a threshold of *p* < 0.05. We then determined the significance of these clusters by comparing their summated t-statistic against a null distribution generated from 5000 random permutations, to identify frequency bands in which information flow was significantly directional.

## Author contribution

M.G., K.I.R., and M.E.D. designed the experiment, M.G., K.I.R., A.O., F.P., and T.P. carried out the data collection. M.G. and F.P. preprocessed the raw electrophysiological data. M.G. analyzed the data. M.G. and M.E.D. interpreted and discussed the results. M.G. and M.E.D. wrote the paper. M.E.D. performed the acquisition of fundings.

## Acknowledgments

We acknowledge the financial support of the the Human Frontier Science Program (http://www.hfsp.org; project RGP0017/2021), the financial support of the Italian Ministry PRIN 2022 funding (contract 20224FWF2J), the financial support of the Italian Ministry PNRR funding (contract P20229752W), the financial support of the Regional Laboratory for Advanced Mechatronics, LAMA FVG (http://lamafvg.it), and the financial support of the European Union Horizon 2020 MSCA Programme (grant 813713). We thank Davide Giana for helpful assistance during behavioral training, surgical and histological procedures. We also thank Marco Gigante, Giulio Iaconcig, and Fabrizio Manzino (CyNexo) for the invaluable technical assistance for hardware construction and software development.

## Supplementary Material

### Single-stimulus trials

The training protocol of the standard task and control task contained *single-stimulus trials*: within a training session under the standard task regime, approximately 20% of trials was constituted by single-stimulus trials, in which only the relevant stimulus was delivered. When rats were trained under the control task, the single-stimulus trials were approximately 40%, within a training session.

**Figure S1:**
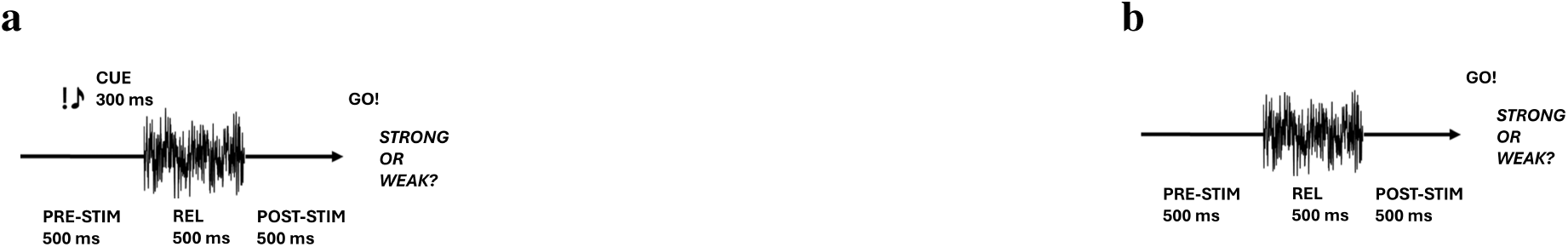
Single-stimulus trials. **a.** Single-stimulus trials in the standard task. **b.** Single-stimulus trials in the control task.

Single-stimulus trials were introduced to ensure that rats stayed in the nosepoke also during the presentation of the first stimulus, and to ensure that they perceived the first stimulus (which corresponded to the irrelevant one, in the case of the standard task and control.

Average performance was lower during single-stimulus trials, for both standard task (Figure S2) and control task (Figure S3). This can be explained by the fact that rats assessed relevance by stimulus position: they learned that the relevant stimulus is the second stimulus of a sequence of two, and when asked to judge the only stimulus of a trial, they might have mistaken REL for IRR.

Despite lower performance, the steepness of the psychometric curve suggest that rats judged REL even when presented alone, excluding the the hypothesis that they entered the nose poke and ignored all events for a fixed period of time.

**Figure S2:**
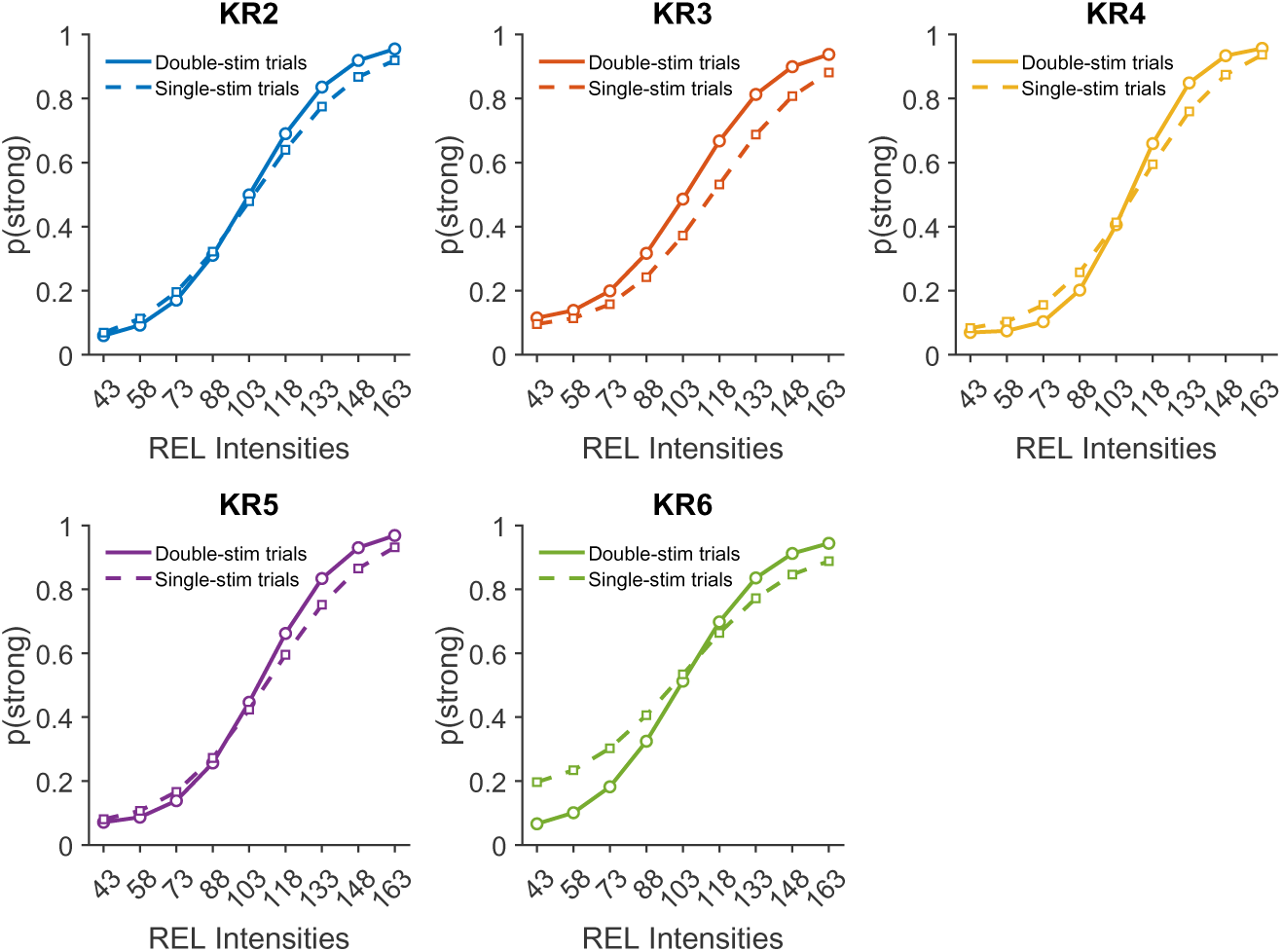
Single-stimulus trials. Single-stimulus trials in the standard task.

**Figure S3:**
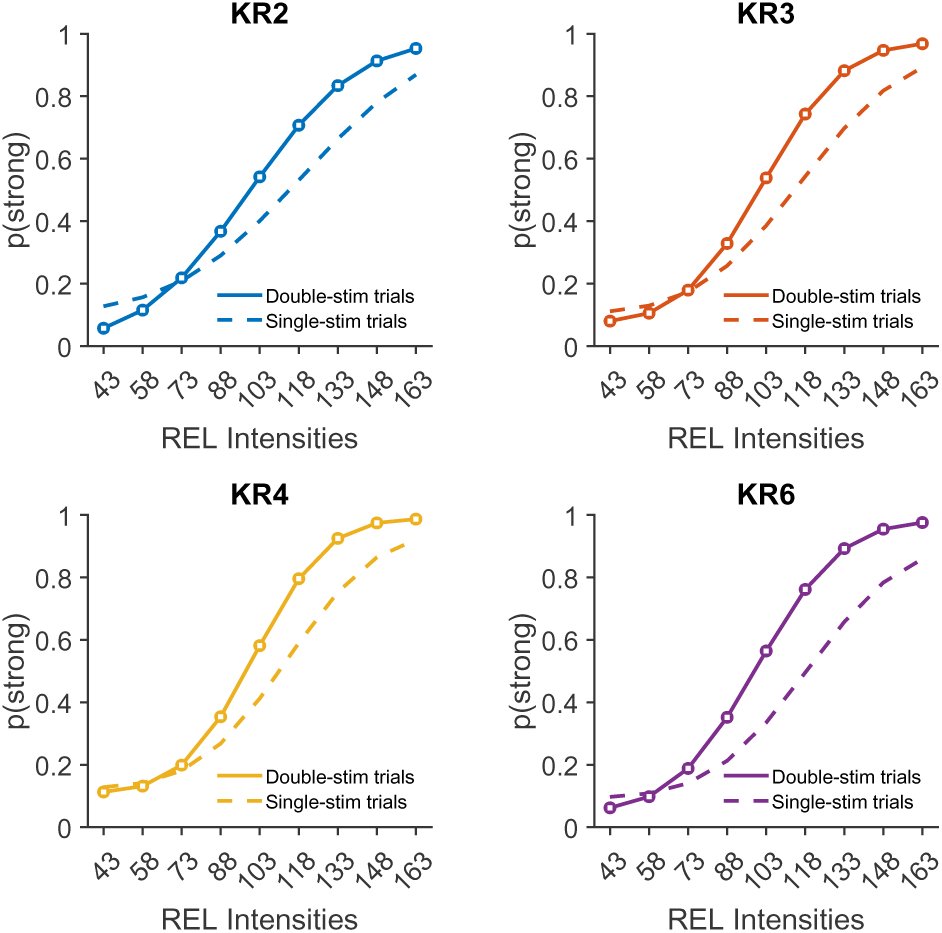
Single-stimulus trials. Single-stimulus trials in the control task.

### Rats trained in the dynamic task do not improve over training

**Figure S4:**
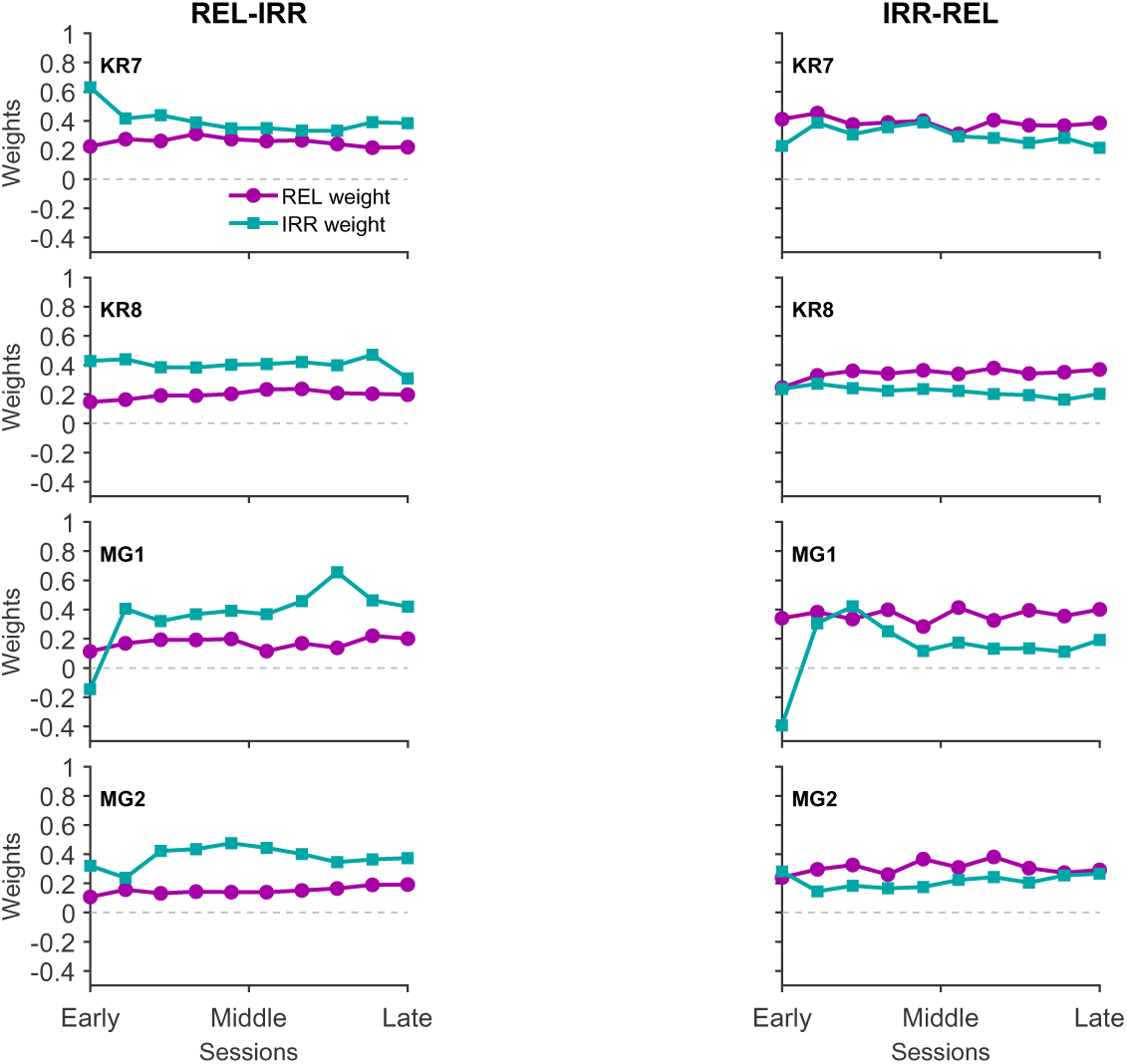
Learning trajectories (dynamic task). Contribution of REL and IRR stimuli to the choice, as described by the GLM weights, for REL-IRR trials (left) and IRR-REL trials (right). All training sessions were binned in 10 groups, and a GLM was computed for each group.

### Linking neural activity to behavior in expert animals

**Figure S5:**
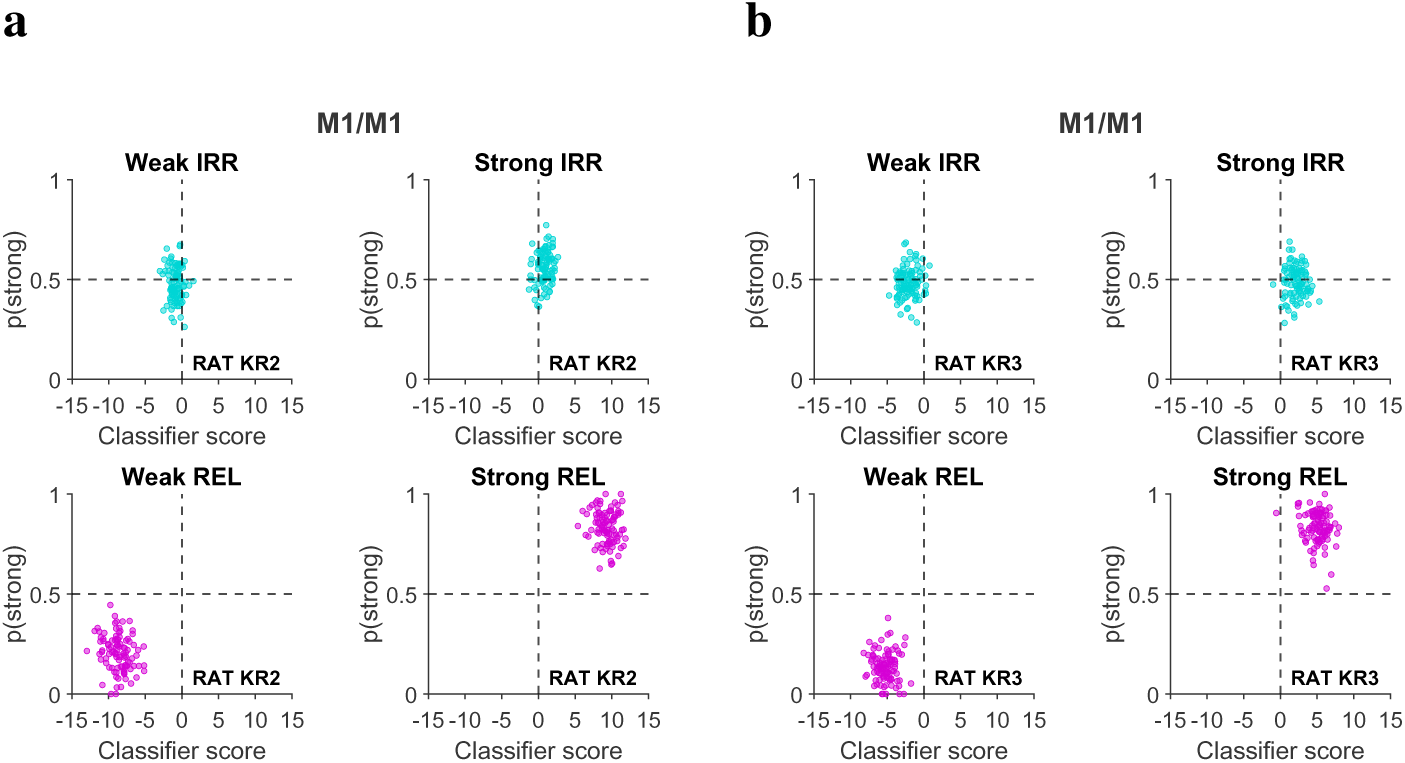
Relationship between decoder’s performance and rat’s choices, for rats KR2 and KR3. **a.** Decoder’s scores plotted against behavioral choices, for each stimulus type, and for each intensity category. Each dot represents the average decoder’s score and rat’s choice for each iteration of the decoder (rat KR2). **b.** Same, for rat KR3.

### Inter-areal coherence

**Figure S6:**
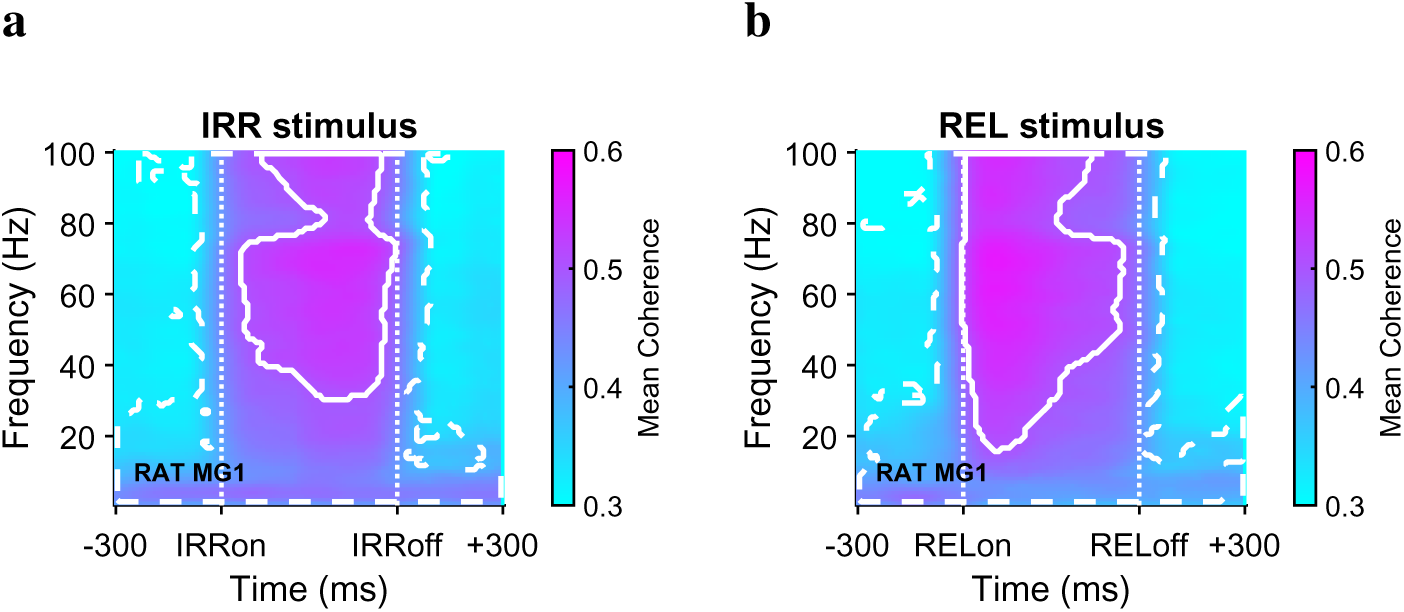
Coherograms during IRR and REL stimulus epochs, for rat MG1. **a.** Coherogram for IRR stimulus epoch (including 300 ms before stimulus onset, and 300 ms after stimulus offset). Dashed lines represent frequency ranges and time bins during which coherence was statistically significant. Solid lines represent frequency ranges and time bins during which coherence was higher than 0.5 **b.** Same, for REL stimulus.

### Frequency domain Granger causality

**Figure S7:**
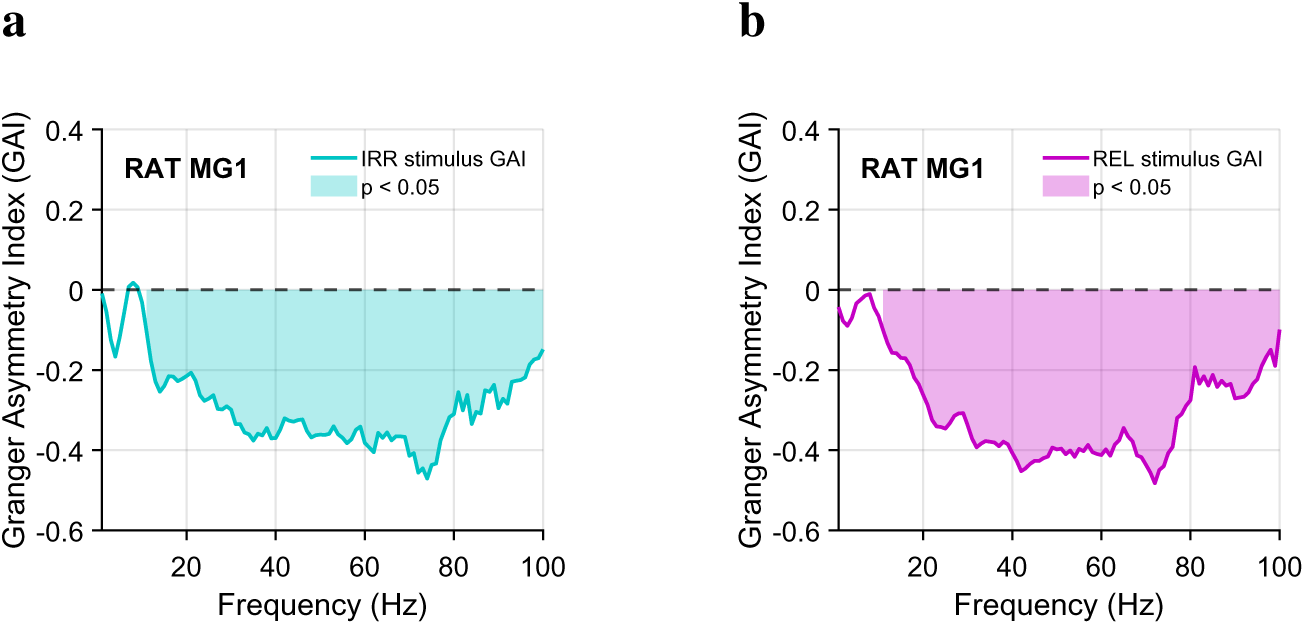
Granger Causality between vS1 and M1/M2, for both REL and IRR epochs, for rat MG1. **a.** Frequency domain Granger Causality during IRR epoch. Shaded areas represent frequencies in which GAI values were statistically significant (*p<0.05*cluster-based permutation test). **b.** Same, but during REL stimulus epoch.

### Time-frequency domain Granger causality

**Figure S8:**
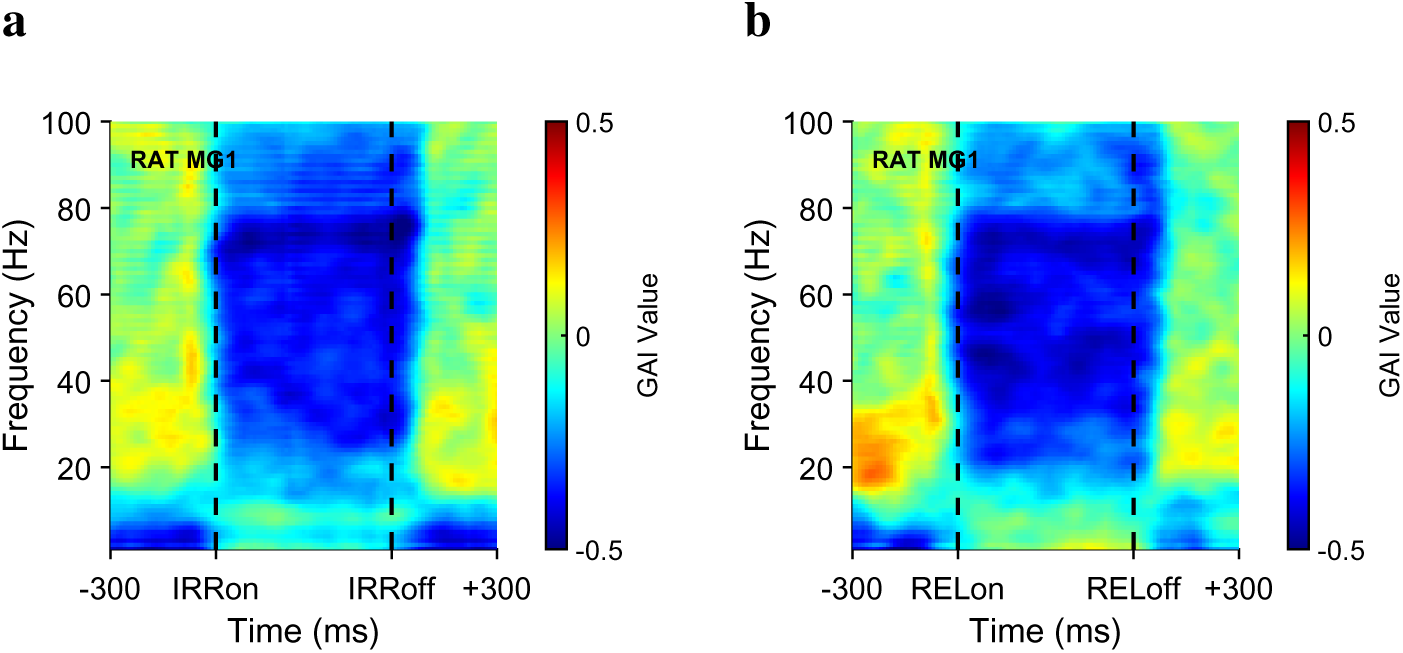
Granger Causality between vS1 and M1/M2, for both REL and IRR epochs, for rat MG1. **a.** Time-frequency domain Granger Causality during IRR stimulus epoch. **b.** Same, for REL stimulus epoch.

**Figure S9:**
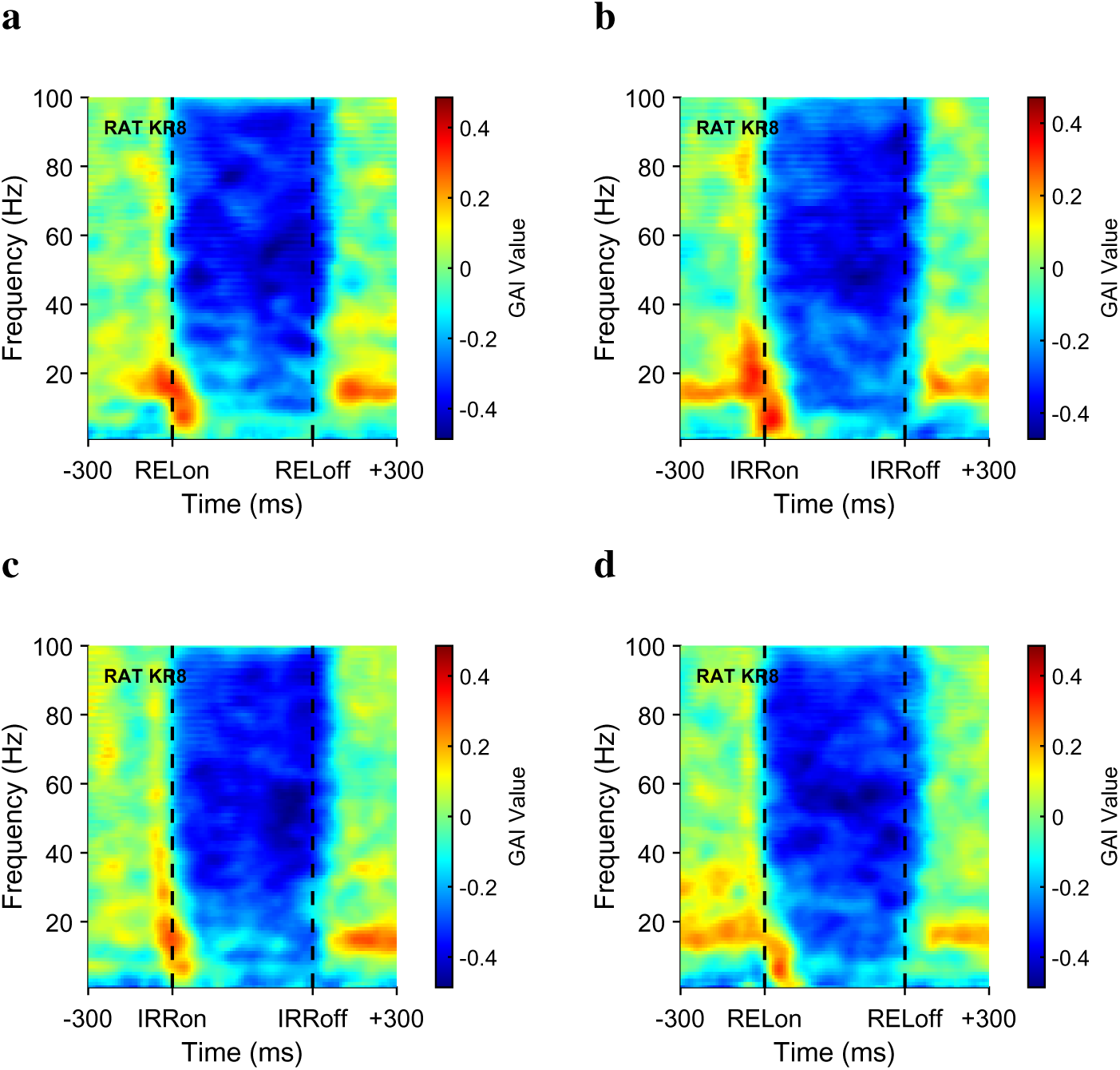
Granger Causality between vS1 and M1/M2, for both REL and IRR epochs, during IRR-REL trials, for rat KR8. **a.** Time-frequency domain Granger Causality during REL stimulus epoch, for REL-IRR trials. **b.** Same, for IRR stimulus. **c.** Time-frequency domain Granger Causality during IRR stimulus epoch, for IRR-REL trials. **d.** Same, for REL stimulus.

